# Precision mRNA Nanomedicine for Targeted Vascular Therapies in ARDS and Atherosclerosis

**DOI:** 10.1101/2025.01.09.632179

**Authors:** Zhengjie Zhou, Jiayu Zhu, Chih-Fan Yeh, Angelo Meliton, David Wu, Ru-Ting Huang, Tzu-Pin Shentu, Aleepta Guha Ray, Destini Wiseman, Erica Budina, Aaron Alpar, Kai-Chien Yang, Jeffrey A Hubbell, Ada Weinstock, Gokhan M Mutlu, Matthew V Tirrell, Yun Fang

**Affiliations:** Department of Medicine, The University of Chicago, IL, USA; Pritzker School of Molecular Engineering, The University of Chicago, IL, USA; Department of Internal Medicine and Cardiovascular Center, National Taiwan University Hospital, Taipei, Taiwan; Graduate Institute and Department of Pharmacology, National Taiwan University College of Medicine, Taipei, Taiwan; Committee on Molecular Medicine, The University of Chicago, IL, USA

## Abstract

Vascular diseases, including acute respiratory distress syndrome (ARDS) and atherosclerosis, are leading causes of global morbidity and mortality, underscoring the critical need for innovative therapies. While human genetics has uncovered specific molecular mechanisms in endothelial cells that drive acute and chronic vascular inflammation central to these conditions, effective *in vivo* strategies to spatiotemporally target these disease-causing pathways using therapeutic mRNAs remain limited. Here, we present a modular and tunable nanoparticle platform (PROGRAMMED nanoparticles) engineered to selectively deliver therapeutic mRNAs to target cells with distinct molecular signatures, such as inflamed endothelial cells expressing vascular cell adhesion molecule 1 (VCAM1). This approach restores key genetics-informed endothelial pathways implicated in disease, including the loss of Krüppel-like factor 2 (KLF2) in acute inflammation underlying ARDS and the suppression of phospholipid phosphatase 3 (PLPP3) in chronic inflammation driving atherosclerosis. In multiple preclinical models, these precision mRNA nanomedicine strategies demonstrated compelling therapeutic efficacy. VCAM1-targeting PROGRAMMED nanoparticles delivering KLF2 mRNA to inflamed lung microvascular endothelial cells significantly improved microvascular health and alleviated virus-induced ARDS. Similarly, PLPP3 mRNA delivery by VCAM1-targeting PROGRAMMED nanoparticles attenuated arterial inflammation, slowed atherosclerosis progression, and promoted regression of advanced plaques. This innovative platform offers a transformative approach for spatiotemporal mRNA therapies targeting inflamed endothelial cells, providing a minimally invasive and highly specific strategy to address acute and chronic vascular diseases. Moreover, the modularity of PROGRAMMED nanoparticles, particularly their ability to display diverse targeting motifs, holds substantial potential for addressing a broad spectrum of vascular complications and other diseases that require precise spatiotemporal correction of dysregulated genes.

## Introduction

Vascular disease is the leading cause of human morbidity and mortality, emphasizing the pressing need for innovative therapeutic strategies. Acute vascular inflammation and loss of capillary integrity in the lung contribute to acute respiratory distress syndrome (ARDS), a major complication for critically ill patients during the influenza and COVID-19 pandemics (*1, 2*). Meanwhile, chronic vascular inflammation and activation are central to atherosclerosis (*3*), the predominant cause of heart attacks and ischemic stroke, which constitute the foremost contributor to global morbidity and mortality among both sexes. Human genetics studies, coupled with mechanistic investigations, have delineated the intricate spatiotemporal dysregulation of key vascular cellular pathways underlying acute and chronic vascular complications (*1, 3*). These pathways represent promising targets for developing novel treatments for conditions such as ARDS or atherosclerosis, which are presently managed predominantly through symptomatic relief measures (e.g., mechanical ventilation and oxygen therapy for ARDS) or by addressing systemic risk factors (e.g., hypercholesterolemia and hypertension for atherosclerosis). However, most of these strategies do not directly intervene in the disease-causing mechanisms within the diseased vasculature itself, emphasizing the necessity for vascular wall-targeted therapeutic innovations.

Messenger RNA (mRNA) therapeutics offer promising avenues for addressing many human diseases, exemplified by the success of mRNA vaccines against COVID-19 (*4, 5*). Precision mRNA therapies targeting specific tissues hold potential for new treatment paradigms by precisely modulating disease-causing molecular pathways within diseased tissues, including the dysfunctional vasculature. However, the effective and targeted delivery of therapeutic mRNA to diseased vascular cells remains a formidable challenge. Vascular endothelial inflammation and dysfunction are key drivers of ARDS and atherosclerosis (*6–8*), necessitating precise targeting of these pathogenic processes for therapeutic intervention. The efficacy of future mRNA-based therapies for ARDS or atherosclerosis hinges on the accurate delivery of functional therapeutic mRNAs to diseased endothelial cells within the lung microvasculature or large elastic and muscular distributing arteries.

Therapeutic targets supported by human genetic evidence of disease have shown a twofold increased likelihood of leading to approved drugs (*9, 10*). Genome-wide association studies (GWAS) and complementary mechanistic investigations have identified tissue-specific roles of the transcription factor Krüppel-like Factor 2 (KLF2) in ARDS (*11, 12*) and the cell-surface glycoprotein phospholipid phosphatase 3 (PLPP3) in atherosclerosis (*13–16*), underscoring their critical contributions to vascular endothelial homeostasis and disease pathogenesis. Acute lung injury induced by viral infection and high-tidal volume ventilation results in the reduction of KLF2, which regulates multiple ARDS GWAS genes related to cytokine storm, oxidation, and coagulation in the lung microvascular endothelium (*11*). Moreover, atherosclerosis predominantly manifests in distinct arterial regions, such as bifurcations and branches, where disturbed blood flow markedly diminishes endothelial PLPP3 levels (*15*). This decrease in PLPP3 expression is exacerbated by the presence of the risk allele at rs17114036 of locus 1p32.2 (*16*), one of the most strongly associated loci with susceptibility to coronary artery disease and ischemic stroke (*13, 14*). The restoration of pulmonary microvascular KLF2 and arterial endothelial PLPP3 is hypothesized to mitigate ARDS and treat atherosclerosis, respectively.

To this end, we have devised an innovative precision mRNA nanomedicine strategy that effectively treats both acute and chronic vascular diseases *in vivo*. We have rationally designed and engineered novel targeted nano-carriers to achieve efficient intracellular delivery of therapeutic mRNAs, tailored to intervene in genetically driven molecular pathways within diseased vascular endothelial cells. These precision mRNA nanomedicine strategies targeting GWAS-linked disease mechanisms have demonstrated significant effectiveness in reducing influenza virus-induced ARDS, slowing atherosclerosis progression, and promoting atherosclerosis regression in mouse models. Our results showcase a potent, scalable, and modular tunable precision mRNA nanomedicine platform capable of spatiotemporally targeting disease-causing molecular mechanisms in specific tissues of interest. These findings offer promising avenues for combating acute and chronic vascular diseases, which pose significant global health and economic challenges.

## RESULTS

### Formulation, optimization, and characterization of lipid nanoparticles for mRNA delivery to vascular endothelial cells

Endothelium-targeting mRNA therapeutics face a significant challenge due to the limited tropism of currently formulated lipid nanoparticles, which impedes efficient delivery of functional mRNA to endothelial cells across various vascular beds within different organs (*17*). In addition, despite the expansive coverage of endothelial cells in the adult human body, vascular diseases typically manifest in localized areas characterized by inflammation. We reasoned that targeted nanoparticle platforms can be engineered to deliver therapeutic mRNAs selectively to inflamed endothelial cells, enhancing the treatment of vascular complications.

Our approach began with the engineering, screening, and selection of leading lipid nanoparticle formulations to achieve effective cytosolic delivery of functional mRNA to vascular endothelial cells. These self-assembled lipid nanoparticles were composed of dioleoylphosphatidylethanolamine (DOPE), lipid-grafted polyamidoamine (PAMAM) dendrimer (G0-C14), cholesterol, 1,2-distearoyl-sn-glycero-3-phosphoethanolamine-poly(ethylene glycol) (DSPE-PEG), and mRNA (Fig 1A). We chose DOPE for its ability to stabilize particles and facilitate endosomal escape by adopting an inverted hexagonal H(II) phase, which destabilizes endosomal membranes (*18–20*). Additionally, we incorporated PAMAM, a highly branched and amine-rich dendrimer, for efficient complexation with negatively charged mRNA through hydrophobic aggregation in aqueous solutions (*21–23*). Cholesterol was included to stabilize the lipid bilayer structure and facilitate the formation of a liquid-ordered phase (*20*), while DSPE-PEG served to increase stability and enhance circulation half-life (*24, 25*). Furthermore, DSPE-PEG can be utilized to conjugate specific ligands to the particle for targeted delivery to desired cells (*26, 27*). We termed this class of structures **PROGRAMMED** nanoparticles (NP) representing “***<u>P</u>****recision **<u>R</u>**NA delivery using cholester**<u>O</u>**l, lipid-**<u>G</u>r**afted P**<u>AM</u>**A**<u>M</u>**, DOP**<u>E</u>**, and **<u>D</u>**SPE-PEG*” (Fig. 1A).

**Figure 1.**
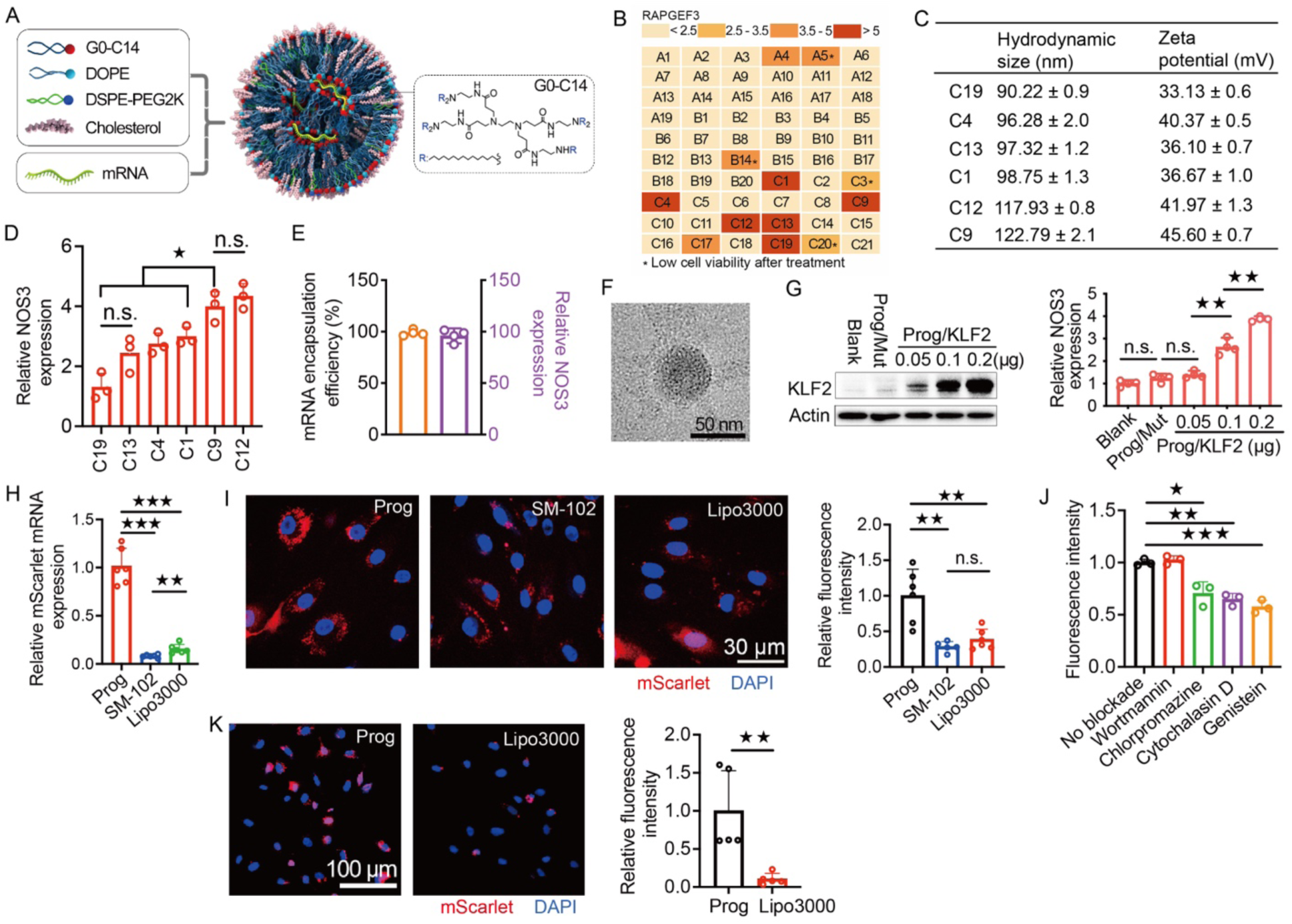
Formulation, selection, and characterization of the PROGRAMMED nanoparticle (Prog) for effective delivery of functional mRNAs to vascular endothelial cells. **(A)** The illustration of self-assembled lipid nanoparticles composed of dioleoylphosphatidylethanolamine (DOPE), lipid-grafted polyamidoamine (PAMAM) dendrimer (G0-C14), cholesterol, 1,2-distearoyl-sn-glycero-3-phosphoethanolamine-poly(ethylene glycol) (DSPE-PEG), and mRNA. **(B)** Gene expression heat map illustrating RAPGEF3 mRNA levels, a key downstream target of KLF2, in human pulmonary microvascular endothelial cells (HPMVEC) 6 hours after the delivery of KLF2 mRNA (200 ng/well in a 24-well plate). Delivery was achieved using nanoparticles formulated with varying molecular ratios of DOPE, PAMAM, G0-C14, cholesterol, and DEPE-PEG. Six KLF2 mRNA-encapsulated nanoparticle formulations induced a greater than five-fold upregulation of RAPGEF3 mRNA expression in HPMVEC. Control nanoparticles with identical chemical formulations but encapsulating untranslatable KLF2 mRNAs containing mutations in all start codons (AUG) were used as negative controls. **(C)** Hydrodynamic size (in nanometers) and zeta potential (in millivolts) of six selected nanoparticles encapsulating KLF2 mRNA, measured using dynamic light scattering. **(D)** mRNA expression levels of Nitric Oxide Synthase 3 (NOS3) in human pulmonary microvascular endothelial cells (HPMVEC) 6 hours after the delivery of KLF2 mRNA (200 ng/well of a 24-well plate) using selected nanoparticles. N = 3. Student’s t-test. **(E)** PROGRAMMED nanoparticles (Prog) consistently demonstrate high efficiency in encapsulating KLF2 mRNA across multiple preparation batches. Moreover, KLF2 mRNA-encapsulated Prog reliably induces NOS3 mRNA expression in HPMVECs across multiple preparation batches. N = 4. **(F)** A negatively stained TEM image of KLF2 mRNA-encapsulated PROGRAMMED nanoparticles. **(G)** PROGRAMMED nanoparticles (Prog) exhibit dose-dependent effects (0.05–0.2 µg/well in a 24-well plate) in delivering functional KLF2 mRNA and inducing the upregulation of NOS3 expression in human pulmonary microvascular endothelial cells (HPMVEC) 6 hours post-delivery. Control nanoparticles encapsulating mutant, non-translatable KLF2 mRNA (Prog/Mut) were used as negative controls. N = 4. Student’s t-test. **(H and I)** PROGRAMMED nanoparticles (Prog) exhibit significantly greater efficacy in delivering functional mScarlet mRNA (200 ng/well in a 24-well plate, assessed 6 hours post-delivery) to pulmonary microvascular endothelial cells (PMVECs) compared to Lipofectamine 3000 and the SM-102-based lipid nanoparticle (LNP) formulation used in the mRNA-1273 COVID-19 vaccine. This enhanced delivery is demonstrated by quantitative real-time PCR analysis of mScarlet mRNA levels (N = 6, one-way Student’s t-test) and fluorescence imaging of mScarlet protein expression (N = 5–6, Student’s t-test). **(J)** Genistein, Cytochalasin D, or Chlorpromazine, each tested separately, significantly reduces the delivery of functional mScarlet mRNA by PROGRAMMED nanoparticles (Prog) in pulmonary microvascular endothelial cells (PMVEC), as demonstrated by decreased mScarlet fluorescence intensity (N = 3, Student’s t-test). **(K)** PROGRAMMED nanoparticles (Prog) demonstrate significantly greater efficiency in delivering functional mScarlet mRNA (200 ng/well in a 24-well plate, assessed 6 hours post-delivery) to human aortic endothelial cells (HAECs) compared to Lipofectamine 3000. This enhanced delivery is quantified through fluorescence imaging analysis of mScarlet protein expression. N = 5, Student’s t-test. All data are represented as mean ± SD. *P < 0.05, **P < 0.01, ***P < 0.001.

A library of sixty PROGRAMMED nanoparticles composed with different molar ratios of the components was formulated and screened for their capacity to deliver functional mRNA effectively to endothelial cells (Table. S1). In human pulmonary microvascular endothelial cells (PMVECs), Rap guanine nucleotide exchange factor 3 (RAPGEF3) is one of the most responsive direct downstream targets transcriptionally up-regulated by KLF2. We measured RAPGEF3 mRNA levels as a surrogate marker of functional KLF2 mRNA delivery. Among the sixty nanoparticles tested (200 ng RNA in a 24-well plate, 6-hour incubation), six exhibited a more than 5-fold upregulation of RAPGEF3 compared to control nanoparticles (Fig. 1B). The control nanoparticles had identical chemical formulations but encapsulated KLF2 mRNAs with mutations in all start codons (AUG), making them untranslatable. These six nanoparticles come in a consistent size range of 90 to 122 nm in diameter, with zeta potentials ranging from 33 to 45 mV (Fig. 1C).

We further prioritized the C9 and C12 PROGRAMMED nanoparticles for study since they are more effective than the rest at increasing the mRNA expression of nitric oxide synthase 3 (NOS3) (Fig. 1D), another key transcriptional target of endothelial KLF2 (*28, 29*). The C9 PROGRAMMED nanoparticle was selected for further characterization and functionalization processes since the expression of the inflammatory markers IL6 and CCL2 is significantly lower in C9-treated PMVECs, compared to C12-treated cells (fig. S1). Consistent mRNA encapsulation (98.8 %) of KLF2 mRNA by the C9 PROGRAMMED nanoparticle and its activity to up-regulate NOS3 in PMVECs by different batches of nanoparticles were demonstrated (Fig. 1E). C9 PROGRAMMED nanoparticle was then characterized with transmission electron microscopy (TEM) and showed around 50 nm in diameter in dry condition (Fig. 1F). Dose-dependent delivery of functional KLF2 mRNA to endothelial cells was detected, accompanied by increased NOS3 expression as a function of elevated KLF2 overexpression; PROGRAMMED nanoparticles encapsulating mutant, non-translatable KLF2 mRNA (Prog/Mut) were used as controls (Fig. 1G). Real-time PCR (Fig. 1H) in PMVECs detected significantly higher mScarlet mRNA when mScarlet mRNA was delivered by C9 PROGRAMMED nanoparticles compared to lipofectamine 3000 and the lipid nanoparticle SM-102 formulation used in the mRNA-1273 COVID-19 vaccine (*30*). The endothelial tropism of C9 PROGRAMMED nanoparticle was demonstrated by their superior effectiveness in mScarlet mRNA delivery, compared to lipofectamine 3000, to PMVECs (Fig. 1I). Functional delivery of mScarlet mRNA to PMVECs was significantly inhibited by genistein, cytochalasin D, and chlorpromazine, but not by wortmannin (Fig. 1J), suggesting a role of caveolae-mediated endocytosis in PROGRAMMED nanoparticle uptake but not micropinocytosis and phagocytosis. C9 PROGRAMMED nanoparticle consistently exhibits potent mRNA delivery efficacy on endothelial cells, including primary human aortic vascular endothelial cells isolated from large arteries, demonstrated by significantly increased mScarlet signals compared to lipofectamine 3000 (Fig. 1K).

### VCAM1-targeting PROGRAMMED NP effectively delivers functional mRNA to inflamed endothelium *in vitro* and *in vivo*

We functionalized the C9 PROGRAMMED NP to deliver therapeutic mRNAs actively to inflamed endothelial cells, a cellular hallmark of vascular diseases such as ARDS and atherosclerosis. Lung inflammation and atherogenic factors (e.g., disturbed flow and hypercholesterolemia) up-regulate endothelial expression of vascular cell adhesion molecule 1 (VCAM-1), the expression of which remains low in healthy blood vessels (*31, 32*). Targeted delivery of therapeutic mRNA to VCAM1-expressing endothelial cells is therefore a potent strategy for ARDS and atherosclerosis therapeutics (*33*). The VCAM1-targeting peptide VHPKQHR, identified by phage display (*34*), was incorporated into the DSPE-PEG molecule (DSPE-PEG-VHPKQHR in Fig. 2A). TEM images showed monodisperse VCAM1-targeting PROGRAMMED NP with a diameter of ∼70 nm in dry conditions and a median hydrodynamic diameter of ∼120 nm in deionized (DI) water (Figs. 2B and C).

**Figure 2.**
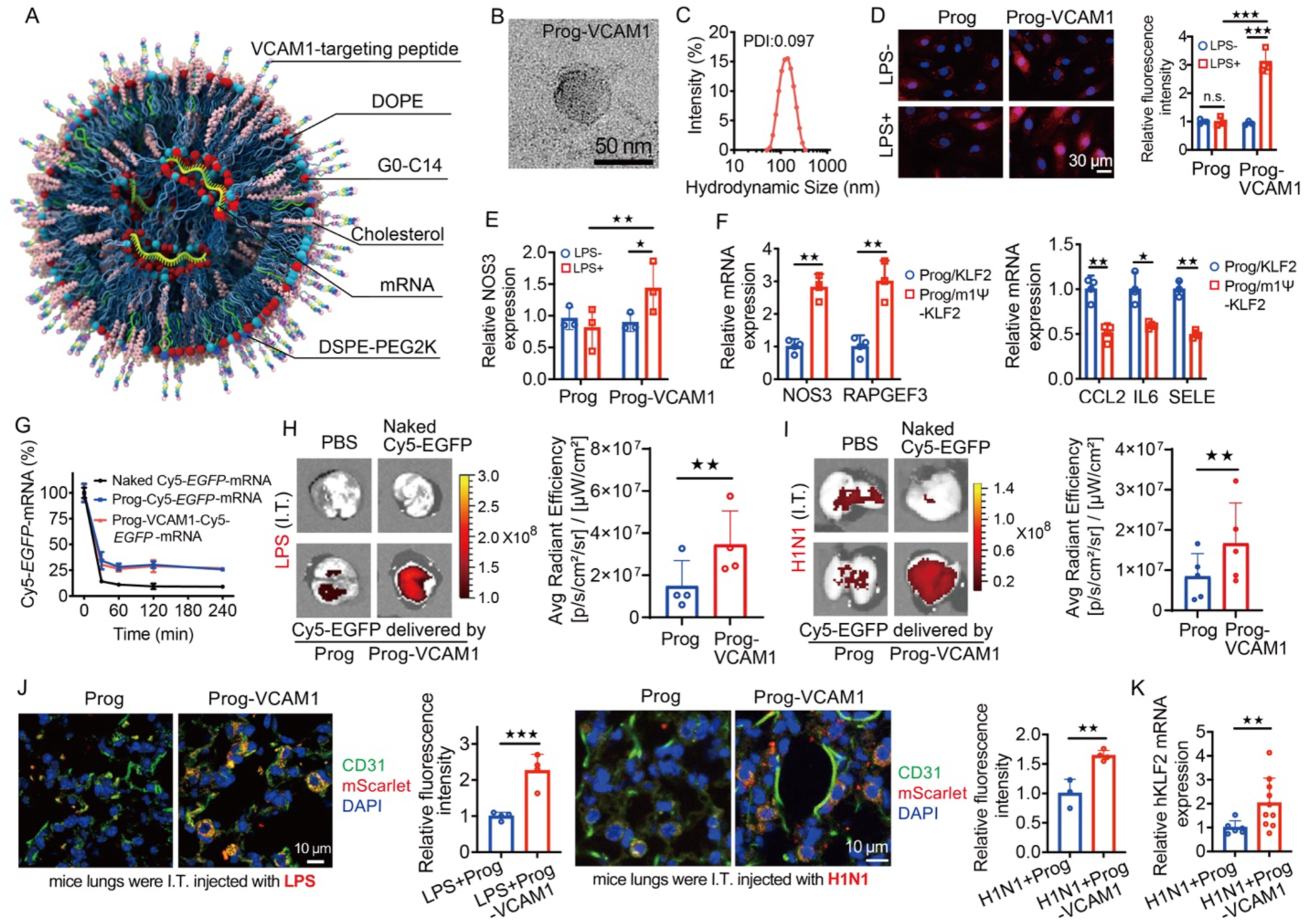
VCAM1-targeting PROGRAMMED nanoparticles (Prog-VCAM1) effectively deliver functional mRNA to inflamed endothelial cells in both in vitro and in vivo settings. **(A)** Schematic representation of mRNA-encapsulated VCAM1-targeting PROGRAMMED nanoparticles (Prog-VCAM1). **(B and C)** A negatively stained transmission electron microscopy (TEM) image and dynamic light scattering (DLS) analysis showing the size distribution of mScarlet mRNA-encapsulated VCAM1-targeting PROGRAMMED nanoparticles (Prog-VCAM1). **(D)** Quantification of mScarlet fluorescence signals demonstrate that Prog-VCAM1 nanoparticles significantly enhance the delivery of functional mScarlet mRNA (200 ng/well in a 24-well plate, assessed 6 hours post-delivery) to LPS-stimulated inflamed HPMVECs compared to quiescent cells. Conversely, non-targeting Prog nanoparticles exhibit similar delivery efficiency in both inflamed and quiescent cells. N = 3. Student’s t-test. **(E)** VCAM1-targeting PROGRAMMED nanoparticles (Prog-VCAM1) demonstrate significantly enhanced delivery of functional KLF2 mRNA (200 ng/well in a 24-well plate, assessed 6 hours post-delivery) to LPS-stimulated inflamed HPMVECs compared to quiescent cells, as indicated by elevated NOS3 mRNA expression. Conversely, non-targeting PROGRAMMED nanoparticles (Prog) exhibit comparable delivery efficiency of functional KLF2 mRNA in both inflamed and quiescent cell conditions. N = 3. Student’s t-test. **(F)** The incorporation of N1-methylpseudouridine (m1Ψ) substitution in KLF2 mRNA significantly enhances its functional efficacy when delivered by PROGRAMMED nanoparticles (Prog). This enhancement is demonstrated by elevated mRNA expression levels of NOS3 and RAPGEF3, along with reduced mRNA expression of CCL2, IL6, and E-electin, in HPMVECs treated with PROGRAMMED nanoparticles encapsulating m1Ψ-modified KLF2 mRNA (Prog/m1Ψ-KLF2, 200 ng/well in a 24-well plate, assessed 6 hours post-delivery), compared to cells treated with PROGRAMMED nanoparticles encapsulating unmodified KLF2 mRNA (Prog/KLF2). N=3. Student’s t-test. **(G)** Half-life assessment of Cy5-labeled eGFP RNA (0.8 mg/kg), either in its unencapsulated form or encapsulated within non-targeting PROGRAMMED nanoparticles (Prog) or VCAM1-targeting PROGRAMMED nanoparticles (Prog-VCAM1), was conducted by measuring the Cy5 signal in murine circulation after intravenous administration via the tail vein. N = 3. Student’s t-test. **(H)** VCAM1-targeting PROGRAMMED nanoparticles (Prog-VCAM1), administered intravenously *via* the tail vein, effectively deliver Cy5-labeled eGFP mRNA (0.8 mg/kg, assessed 24 hours post-delivery), to inflamed mouse lungs subjected to intratracheal (IT) LPS administration. This is demonstrated by a significantly increased Cy5 signal, compared to the minimal signal observed with non-targeting PROGRAMMED nanoparticles (Prog). N = 4. Student’s t-test. **(I)** VCAM1-targeting PROGRAMMED nanoparticles (Prog-VCAM1), administered intravenously via the tail vein, effectively deliver Cy5-labeled eGFP mRNA (0.8 mg/kg, evaluated 24 hours post-administration) to inflamed mouse lungs subjected to intratracheal (IT) influenza H1N1 virus administration. This is demonstrated by a significantly higher Cy5 signal compared to the minimal signal observed with non-targeting PROGRAMMED nanoparticles (Prog). N = 5. Student’s t-test. **(J)** Immunofluorescence analysis of lung sections from mice subjected to intratracheal administration of LPS (N = 4, one-way Student’s t-test) or H1N1 virus (N = 3-4, one-way Student’s t-test) reveals a robust mScarlet signal predominantly localized to CD31-positive endothelial cells, 24 hours following intravenous administration of VCAM1-targeting PROGRAMMED nanoparticles (Prog-VCAM1) encapsulating mScarlet mRNA (1 mg/kg). In contrast, lungs treated with non-targeting PROGRAMMED nanoparticles (Prog) exhibit minimal mScarlet signal, highlighting the specificity and efficacy of Prog-VCAM1 in targeting inflamed pulmonary endothelial cells. **(K)** Real-time PCR analysis demonstrates a significant increase in delivered KLF2 transcript levels in H1N1-infected mouse lungs 24 hours following intravenous administration of KLF2 mRNA (0.8 mg/kg) encapsulated in VCAM1-targeting PROGRAMMED nanoparticles (Prog-VCAM1), compared to H1N1-infected mice treated with KLF2 mRNA encapsulated in non-targeting PROGRAMMED nanoparticles (Prog). N = 6-10. Student’s t-test. <u>All data are represented as mean ± SD. *P < 0.05, **P < 0.01, ***P < 0.001.</u>

Lipopolysaccharides (LPS) were used to induce VCAM-1 expression in PMVECs (fig. S2). There was no difference in mScarlet mRNA delivery by non-targeting PROGRAMMED NP in control and LPS-treated cells (Fig. 2D). In contrast, VCAM1-targeting PROGRAMMED NP delivered functional mRNA more effectively to LPS-treated than to quiescent endothelial cells, demonstrated by increased mScarlet signaling resulting from mScarlet mRNA delivery (Fig. 2D), and by elevated NOS3 expression resulting from KLF2 mRNA delivery (Fig. 2E). Incorporation of N1-methylpseudouridine (m1Ψ) substitution of the uridine, an RNA modification associated with enhanced protein expression and reduced immunogenicity (*4, 35*), significantly enhanced the functional output of KLF2 mRNAs delivered by PROGRAMMED NP, demonstrated by increased expression of NOS3 and RAPGEF3, and by reduced expression of IL-6, CCL2, and E-selectin in PMVEC when m1Ψ KLF2 mRNA was used (Fig. 2F). For the remaining studies, we then prioritized the VCAM1-targeting PROGRAMMED NP for the delivery of m1Ψ-modified mRNA.

*In vivo*, both non-targeting and VCAM1-targeting PROGRAMMED NPs enhanced the half-life of Cy5-labeled (eGFP) RNA in mouse circulation after intravenous administration *via* the tail vein (Fig. 2G). In the lungs of healthy adult male C57BL/6J mice, *In Vivo* Imaging System (IVIS) did not detect significant Cy5-labeled mRNA delivered intravenously in naked form or by non-targeting or VCAM1-targeting C9 PROGRAMMED NPs (fig. S3). Lung inflammation was induced in adult male mice through intratracheal delivery of LPS (intratracheal injection of 50 µL (1 mg/mL) LPS or influenza A virus (A/Puerto Rico/8/1934 [H1N1], PR8, 200 PFU/mouse). Intravenous administration of VCAM1-targeting PROGRAMMED NP effectively delivered Cy5-labeled eGFP mRNA, as evidenced by Cy5 signaling, to inflamed lung induced by LPS or H1N1 viruses (Fig. 2H and Fig. 2I). Very low Cy5 signal was detected when Cy5-labeled mRNA was delivered by non-targeting PROGRAMMED NP.

Immunofluorescence images of sections from LPS- or H1N1 virus-treated lung showed significant mScarlet signal, predominantly localized in CD31-positive endothelial cells, when mScarlet mRNA was delivered by VCAM1-targeting PROGRAMMED NP, while no significant mScarlet signal was detected when non-targeting PROGRAMMED NP was employed (Fig. 2J). In agreement with these results, real-time PCR detected a significant increase in delivered KLF2 transcripts in the inflamed mouse lung subjected to H1N1 viruses following intravenous injections of human KLF2 mRNA-encapsulated VCAM1-targeting PROGRAMMED NP, compared to mice subjected to injections of KLF2 mRNA-encapsulated non-targeting PROGRAMMED NP (Fig. 2K). These *in vitro* and *in vivo* results collectively demonstrate that VCAM1-targeting PROGRAMMED NP effectively drives active delivery of functional mRNA to inflamed microvascular endothelial cells in the lung.

The biosafety of VCAM1-targeting PROGRAMMED NP was evaluated using critical biomarkers in circulation and histological sections of major organs. Adult male C57BL/6J mice were intravenously injected with VCAM1-targeting PROGRAMMED NP encapsulating mutant or functional KLF2 mRNA. Blood samples were collected at 24 and 48 hr post-injection to measure the activities of alanine aminotransferase (ALT), aspartate aminotransferase (AST), and amylase, along with the concentrations of blood urea nitrogen (BUN), creatinine, bilirubin, albumin, and total protein (fig. S4). Liver health remained unaffected following administration of VCAM1-targeting PROGRAMMED NP, as evidenced by comparable activities of ALT, AST, and bilirubin levels between PBS- and NP-treated mice. Kidney function was normal, indicated by similar levels of BUN and creatinine, while standard pancreatic function was demonstrated by comparable amylase activity across groups. Additionally, unchanged levels of albumin and total protein further confirmed normal kidney and liver functions in mice treated with VCAM1-targeting PROGRAMMED NP. Histological sections of the heart, lung, spleen, liver, and kidney did not reveal significant differences in these tissues in mice 48 hr post-administration compared to PBS-treated controls (fig. S5). These results collectively suggest a favorable safety profile for the VCAM1-targeting PROGRAMMED NP.

### KLF2 mRNA-encapsulated VCAM1-targeting PROGRAMMED NP significantly lessened ARDS induced by influenza A virus

The therapeutic effectiveness of VCAM1-targeting PROGRAMMED NP for targeted mRNA therapy treating ARDS *in vivo* was determined in mice subjected to intratracheal challenge with PR8 H1N1 influenza A virus. Genetic, animal, and human studies, alongside mechanistic investigations, have collectively shown that the downregulation of the transcription factor KLF2 in inflamed pulmonary microvascular endothelial cells is a prominent molecular hallmark leading to heightened vascular permeability and lung inflammation(*11, 12, 36*), which are characteristic features of ARDS. We employed VCAM1-targeting PROGRAMMED NP to deliver KLF2 mRNA to inflamed lung microvascular endothelial cells. Un-translatable KLF2 mRNAs with mutations of all start codons (AUG) were engineered and used as control RNA (Mut). Adult male C57BL/6J mice were subjected to intratracheal instillation of PR8 H1N1 influenza A virus (200 PFU/mouse) to induce acute lung injury. VCAM1-targeting PROGRAMMED NP carrying wildtype or mutant human KLF2 mRNA were intravenously administered (0.8 mg mRNA/kg body weight, total 100 µL) on day 1 and day 4 after the intratracheal instillation of H1N1 virus (Fig. 3A). Mice were sacrificed on Day 6 (Fig. 3A). H1N1 viruses caused ARDS evidenced by significantly elevated proteins and cell counts in the fluid of bronchoalveolar lavage (BAL) (Fig. 3B), accompanied by increased expression of inflammatory markers IL-6, and TNF-α in the lung BAL (Fig. 3C). H1N1 virus-induced lung injury was significantly lessened by injections of KLF2 mRNA-encapsulated VCAM1-targeting PROGRAMMED NP, demonstrated by reduced cell counts/proteins in BAL fluid and decreased expression of the inflammatory markers (Fig. 3B and 3C). Notably, administrations of VCAM1-targeting PROGRAMMED NP carrying mutant KLF2 mRNA had no therapeutic effect on reducing lung edema and inflammation. The histological scores, graded as follows: 0: no damage; 1: mild; 2: moderate; 3: severe; and 4: very severe injury, with an increment of 0.5 if the lung injury fell between two integers (*37, 38*), were conducted on paraffin-embedded, sectioned lungs to assess and quantify the severity of lung injury. Consistent with the measurements of BAL and gene expression, the lung histological score was significantly increased due to H1N1 virus-induced ARDS, which was not affected by VCAM1-targeting PROGRAMMED NP carrying mutant KLF2 mRNA but was remarkably reduced by the administration of KLF2 mRNA-encapsulated PROGRAMMED NP (Fig. 3D and 3E). These results demonstrated that KLF2 mRNA-encapsulated VCAM1-targeting PROGRAMMED NP effectively lessens virus-induced lung injury *in vivo*.

**Figure 3.**
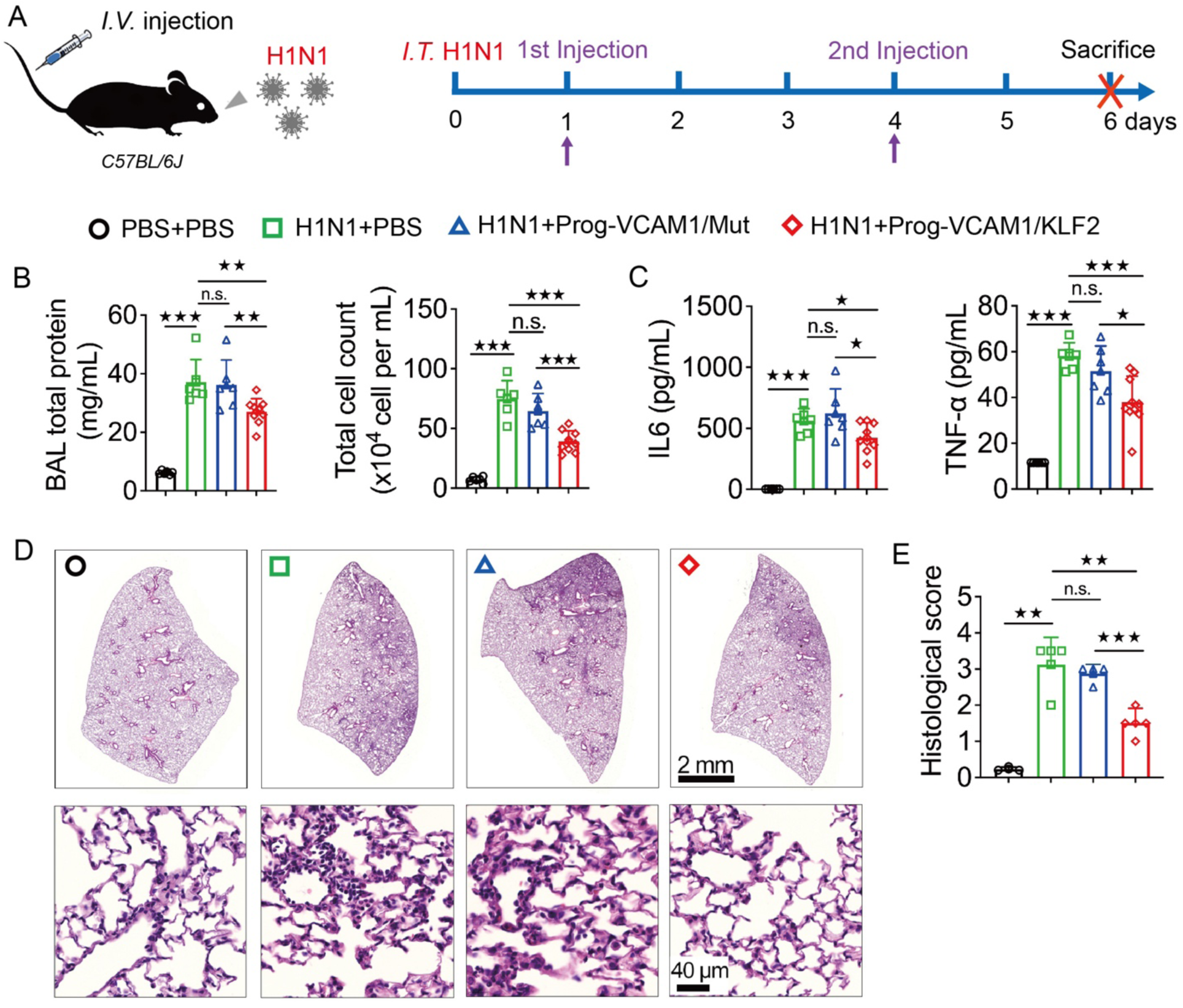
VCAM1-targeting PROGRAMMED nanoparticles encapsulating KLF2 mRNA (Prog-VCAM1/KLF2) effectively alleviate influenza A virus-induced ARDS in a mouse model. **(A)** ARDS was induced in adult C57BL/6J mice through intratracheal instillation of 200 PFU of influenza A virus. VCAM1-targeting PROGRAMMED nanoparticles (Prog-VCAM1), encapsulating either functional or mutant KLF2 mRNA (0.8 mg/kg, 100 µL), were administered intravenously via the tail vein on days 1 and 4 post-instillation. Mice were euthanized on day 6 for analysis. **(B)** H1N1 virus-induced lung injury is significantly mitigated by injections of VCAM1-targeting PROGRAMMED nanoparticles (Prog-VCAM1) encapsulating functional KLF2 mRNA (0.8 mg/kg), as evidenced by reduced cell counts and protein levels in bronchoalveolar lavage (BAL) fluid. In contrast, Prog-VCAM1 carrying non-translatable KLF2 mRNA shows no such effect. N = 6-9. Student’s t-test. **(C)** H1N1 virus-induced lung inflammation is significantly alleviated by the administration of VCAM1-targeting PROGRAMMED nanoparticles (Prog-VCAM1) encapsulating functional KLF2 mRNA (0.8 mg/kg), as evidenced by a marked reduction in the transcriptional expression of inflammatory markers. In contrast, Prog-VCAM1 encapsulating non-translatable KLF2 mRNA does not demonstrate this effect. N = 6-9. Student’s t-test. **(D)** H1N1 virus-induced ARDS, assessed through lung histology, is significantly alleviated by the administration of VCAM1-targeting PROGRAMMED nanoparticles (Prog-VCAM1) encapsulating functional KLF2 mRNA (0.8 mg/kg), while Prog-VCAM1 encapsulating non-translatable KLF2 mRNA exhibits no such effect (representative images shown). N = 6-9. Student’s t-test. **(E)** H1N1 virus-induced ARDS, assessed by lung histological scoring, was significantly reduced following the administration of functional KLF2 mRNA (0.8 mg/kg)-encapsulated VCAM1-targeting PROGRAMMED nanoparticles (Prog-VCAM1). In contrast, Prog-VCAM1 carrying non-translatable KLF2 mRNA showed no therapeutic effect. N= 3-4. Student’s t-test. <u>All data are represented as mean ± SD. *P < 0.05, **P < 0.01, ***P < 0.001.</u>

### PLPP3 mRNA-encapsulated VCAM1-targeting PROGRAMMED NP significantly reduced disturbed flow-induced atherosclerosis

Genome-wide association studies along with mechanistic investigations collectively demonstrated that endothelial PLPP3 reduction drives arterial inflammation and disturbed flow-induced atherosclerosis (*13–16*). PLPP3 restoration in inflamed arterial endothelial cells, although remaining challenging, is an attractive approach to lessen atherogenesis. The endothelial tropism of PROGRAMMED NP was demonstrated by its effectiveness of delivering functional m1Ψ PLPP3 mRNA (200 ng RNA in a 24-well plate, 6-hour incubation) to human aortic endothelial cells *in vitro*, showing expression of Flag-tagged PLPP3 in human aortic endothelial cells (HAEC) (Fig. 4A), accompanied by reduced expression of inflammatory genes *IL6*, *CCL2*, and *SELE* (Fig. 4B). In addition, VCAM1-targeting functionalization significantly enhanced the delivery of functional mScarlet mRNA to inflamed HAECs induced by LPS (fig. S6).

**Figure 4.**
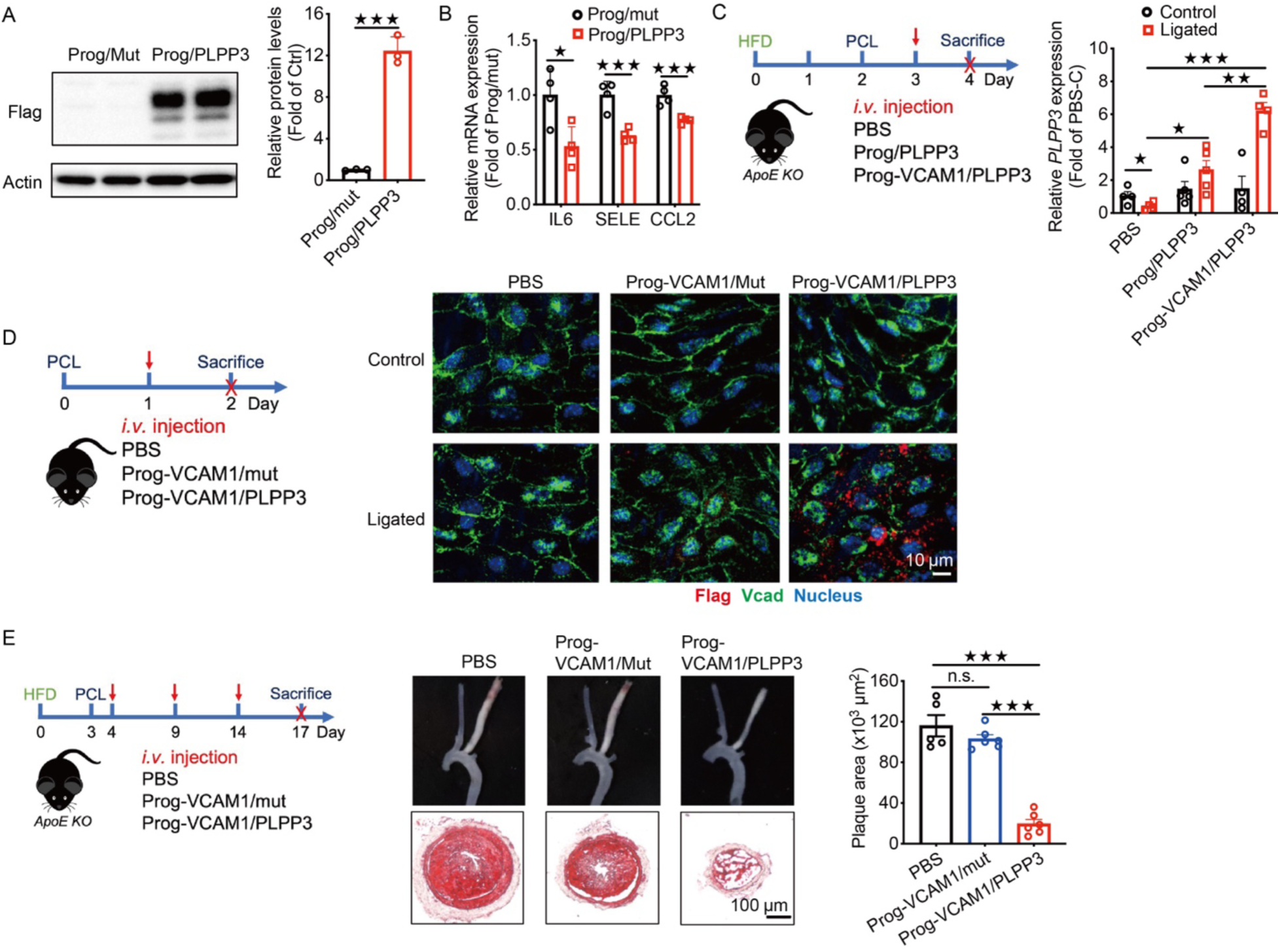
VCAM1-targeting PROGRAMMED nanoparticles encapsulating PLPP3 mRNA (Prog-VCAM1/PLPP3) significantly attenuate disturbed flow-induced atherosclerosis in a mouse model. **(A)** PROGRAMMED nanoparticles (Prog/PLPP3) effectively deliver functional m1Ψ-modified mRNA encoding FLAG-tagged PLPP3 (200 ng/well in a 24-well plate) to cultured human aortic endothelial cells (HAEC), as demonstrated by increased FLAG-tagged PLPP3 protein levels in Western blot analysis. In contrast, control PROGRAMMED nanoparticles (Prog/mut), encapsulating untranslatable PLPP3 mRNA with mutations in all start codons (AUG), show no detectable effect. N = 3. Student’s t-test. **(B)** Transcriptional expression of IL6, SELE, and MCP1 is significantly reduced in HAEC treated with PLPP3 mRNA (200 ng/well in a 24-well plate) delivered by PROGRAMMED nanoparticles (Prog/PLPP3), while no such reduction is observed with PROGRAMMED nanoparticles (Prog/mut) encapsulating untranslatable PLPP3 mRNA (0.8 mg/kg), as determined by real-time PCR analysis. n=4. Student’s t-test. **(C)** PLPP3 mRNA levels in the endothelium-enriched intima were significantly reduced in the partially ligated left carotid artery (LCA) compared to the non-ligated right carotid artery (RCA) in *Apoe*^−/−^ mice. Intravenous administration of PLPP3 mRNA (0.8 mg/kg)-encapsulated PROGRAMMED nanoparticles, either non-targeting (Prog) or VCAM1-targeting (Prog-VCAM1), significantly restored PLPP3 mRNA levels in the intima of the ligated LCA. Importantly, Prog-VCAM1 achieved markedly greater PLPP3 mRNA restoration in endothelial cells of the ligated LCA compared to Prog. n=4-5. Student’s t-test. **(D)** *En face* immunofluorescence analysis, performed one day following intravenous administration of PLPP3 mRNA (2 mg/kg)-encapsulated VCAM1-targeting PROGRAMMED nanoparticles (Prog-VCAM1), demonstrates the presence of lag-tagged PLPP3 protein localized to the endothelium of the ligated left carotid artery (LCA) in *Apoe*^−/−^ mice. In contrast, no lag-tagged PLPP3 protein is detected in the non-ligated right carotid artery (RCA) or in the ligated LCA of mice treated with Prog-VCAM1 encapsulating non-translatable PLPP3 mRNA containing mutations in all start codons (AUG). **(E)** Disturbed flow-induced atherosclerosis in the ligated left carotid artery (LCA) of high-fat diet-fed *Apoe^−/−^* mice is significantly reduced following intravenous administration of VCAM1-targeting PROGRAMMED nanoparticles (Prog-VCAM1) encapsulating functional PLPP3 mRNA (0.8 mg/kg, three injections). In contrast, Prog-VCAM1 encapsulating non-translatable mutant PLPP3 mRNA exhibits no effect on carotid lesion size, underscoring the therapeutic specificity of functional PLPP3 mRNA delivery. n=5-6. Tukey’s post hoc test after one-way ANOVA. <u>(A) and (B) are represented as mean ± SD, (C) and (E) are represented as</u> <u>mean ± SEM. *P < 0.05, **P < 0.01, ***P < 0.001.</u>

We therefore engineered the VCAM1-targeting PROGRAMMED NP to encapsulate m1Ψ human PLPP3 mRNA, aiming to drive active delivery of PLPP3 mRNA to inflamed arterial endothelial cells and lessen atherosclerosis. Partial carotid artery ligation (fig. S8) was conducted in the left carotid artery (LCA) of high fat-fed *Apoe^−/−^* mice to induce acute disturbed flow leading to induced VCAM-1 and reduced PLPP3 expression in endothelial cells, when compared to those in the non-ligated right carotid artery (RCA) (*15*). A single intravenous injection of PLPP3 mRNA-encapsulated, VCAM1-targeting PROGRAMMED nanoparticles (0.8 mg mRNA/kg body weight, total 100 µL) *via* the tail vein significantly restored and enhanced endothelial PLPP3 expression in the ligated LCA, without affecting quiescent endothelial cells in the non-ligated RCA (Fig. 4C). Un-translatable Flag-tagged PLPP3 mRNA with mutations of all start codons (AUG) were engineered and used as control RNAs. *En face* immunofluorescence images one day after the injection of VCAM1-targeting PROGRAMMED NP (2 mg mRNA/kg body weight, total 100 µL) detected the tagged PLPP3 expression in the ligated LCA, but not the non-ligated RCA, in male *Apoe^−/−^*mice, demonstrating active delivery of functional mRNA to inflamed endothelial cells *in vivo* (Fig. 4D). No Flag-PLPP3 protein was detected in mice subjected to control mutant PLPP3 mRNA delivered by the VCAM1-targeting PROGRAMMED NP. In agreement with these results, the VCAM1-targeting PROGRAMMED NP, but not the non-targeting NP, effectively delivered functional mScarlet mRNA (2 mg mRNA/kg body) to inflamed endothelial cells in the ligated carotid artery in mice (fig. S8).

The therapeutic effectiveness of the VCAM1-targeting PROGRAMMED NP in lessening atherosclerosis progression was tested *in vivo*. The study design is illustrated in Fig. 4E, in which disturbed blood flow-induced atherosclerosis was shown by Oil red O staining in the ligated LCA 14 days after the surgery, while the RCA developed no sign of vascular remodeling in the high fat-fed male *Apoe^−/−^* mice. Functional or mutant control PLPP3 mRNA, encapsulated in VCAM1-targeting PROGRAMMED NP (0.8 mg mRNA/kg body weight, total 100 µL), was intravenously administered on days 1, 6, and 11 after the surgery. Disturbed flow-induced carotid lesions were markedly reduced (by 83%) by the treatments of functional PLPP3 mRNA-encapsulated VCAM1-targeting PROGRAMMED NP when compared to PBS-injected controls. In sharp contrast, injections of VCAM1-targeting PROGRAMMED NP encapsulating mutant PLPP3 mRNA had no effect on the size of carotid lesions. Body weights and plasma cholesterol levels were not altered by the injections of mRNA-encapsulated VCAM1-targeting PROGRAMMED NP (fig. S9). These data collectively demonstrated that VCAM1-targeting PROGRAMMED NP effectively drives active mRNA delivery to inflamed arterial endothelial cells and endothelial PLPP3 restoration by targeted mRNA therapy significantly reduces progression of atherosclerosis *in vivo*.

### PLPP3 mRNA-encapsulated VCAM1-targeting PROGRAMMED NP significantly promotes regression of advanced atherosclerotic plaques

The therapeutic efficacy of VCAM1-targeting PROGRAMMED NP in treating chronic vascular complications was evaluated by examining the potential of m1Ψ PLPP3 mRNA-encapsulated NP to promote regression of advanced atherosclerotic plaques in mice. The transformation of inflamed atherosclerotic lesions into a more benign fibrotic scar-like histology, often termed plaque regression, is pivotal in the development of atherosclerosis therapy for cardiovascular patients due to its potential to stabilize plaques and reduce major adverse cardiovascular events such as heart attacks and ischemic strokes (*39, 40*). Indeed, most patients with atherosclerotic cardiovascular disease are undergoing treatment with potent cholesterol-reducing medications and/or lifestyle changes. In humans, aggressive cholesterol lowering is associated with decreased plaque inflammation, an increase in fibrous cap thickness, and reduced major adverse cardiovascular events (*39–41*). We hypothesize that PLPP3 mRNA-encapsulated VCAM1-targeting PROGRAMMED NP is an effective vascular wall-targeted therapeutic approach to promote further lesion regression in subjects undergoing aggressive cholesterol lowering through pharmacological treatments and lifestyle changes. To test this, advanced atherosclerosis was induced in male C57BL/6 mice through the administration of adeno-associated virus 9 expressing proprotein convertase subtilisin/kexin type 9 PCSK9 (AAV9-PCSK9) along with a high-fat diet (HFD) (Fig. 5A). Mice were sacrificed 4 months after viral transduction and initiation of the high-fat diet (baseline group), revealing significantly elevated plasma cholesterol levels (Fig. 5B) and advanced plaques (Fig. 5C) in the aortic roots characterized by lipids and CD68+ monocytes/macrophages. To simulate lesion regression similar to that observed in humans undergoing aggressive cholesterol-lowering therapy, hypercholesterolemic mice were switched to a non-high-fat chow diet supplemented with the microsomal triglyceride transfer protein (MTP) inhibitor Lomitapide for 2 weeks prior to harvesting (Fig. 5A). This approach has been shown to rapidly reduce plasma lipid levels and induce plaque regression, as evidenced by a significant decrease in macrophages (CD68+) and an increase in collagen content (*42*)—a marker associated with enhanced plaque stability in human atherosclerosis. In AAV9-PCSK9-injected mice maintained on a high-fat diet (HFD) for 4 months, transitioning to a chow diet supplemented with an MTP inhibitor led to a marked reduction in plasma cholesterol levels, decreasing from 1036 mg/dL to ∼160 mg/dL within two weeks (regression group) (Fig. 5B). These findings are consistent with a previous study demonstrating that transitioning to a chow diet, coupled with treatment using an MTP inhibitor, effectively normalized hyperlipidemia (*43*). During the two-week lesion regression period, PBS or VCAM1-targeting PROGRAMMED nanoparticles encapsulating either mutant PLPP3 mRNA or functional PLPP3 mRNA (0.8 mg mRNA/kg body weight, administered in three injections on days 1, 6, and 11 following the chow diet and MTP inhibitor treatment) were delivered intravenously *via* the tail vein (Fig. 5A). . In mice subjected to cholesterol reduction and intravenous administration of PBS, the lesion size remained comparable to those observed before cholesterol reduction (Fig. 5C and 5D). However, cholesterol reduction significantly decreased the accumulation of CD68+ macrophages in plaques, as indicated by the total CD68+ lesion area (Fig. 5C and fig. S10) and the percentage of CD68+ area within the lesions (Fig. 5C and 5E). Additionally, cholesterol reduction resulted in increased collagen content within the lesions (Fig. 5C and 5F). These findings collectively demonstrate regression of advanced plaques, characterized by reduced inflammation and increased collagen content, in mice as a result of the pharmacological intervention and diet switch.

**Figure 5.**
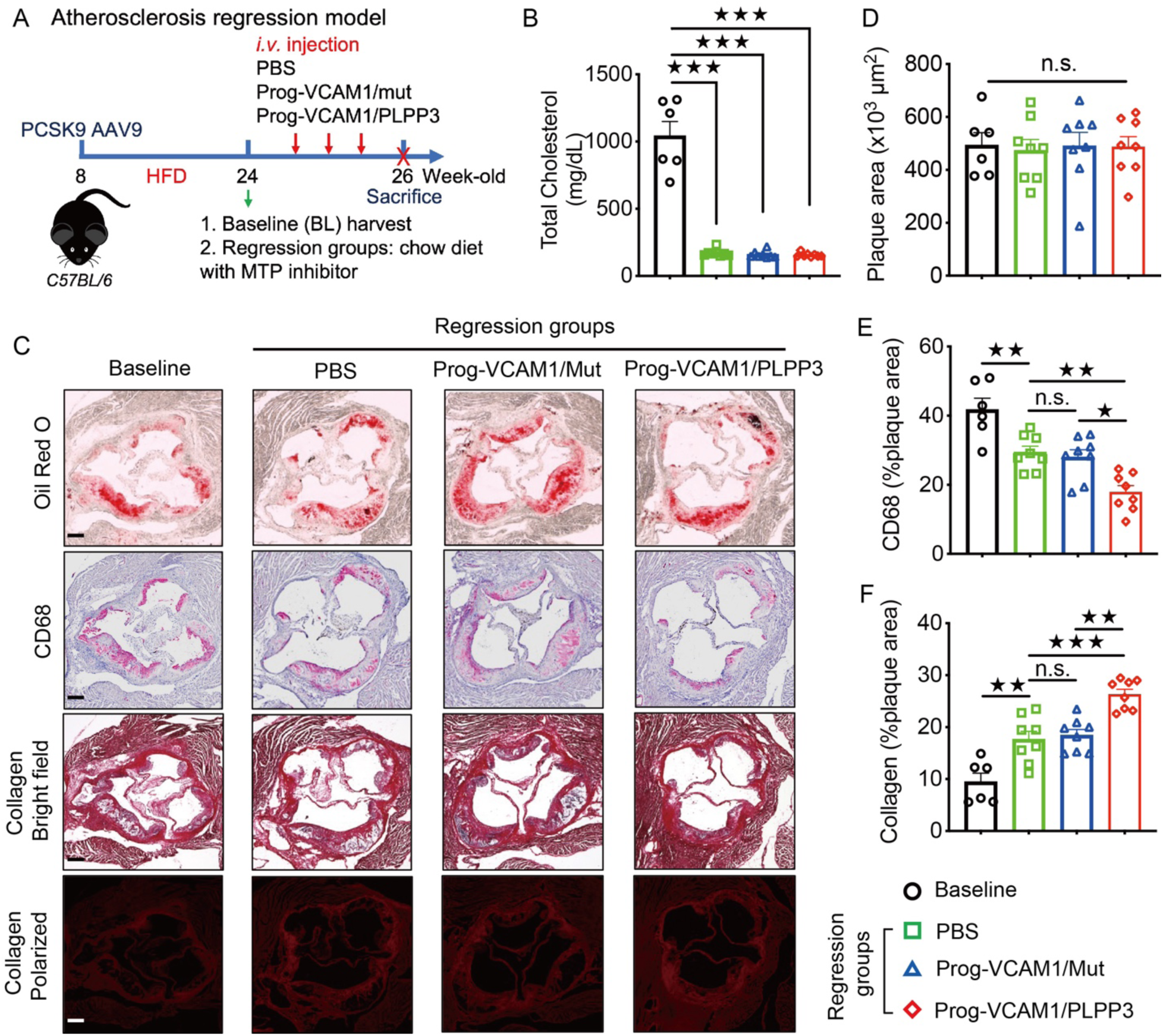
PLPP3 mRNA-encapsulated VCAM1-targeting PROGRAMMED nanoparticles (Prog-VCAM1/PLPP3) significantly promote the regression of advanced atherosclerotic plaques. **(A)** Advanced atherosclerosis was induced in C57BL/6 mice by AAV9-PCSK9 administration combined with a high-fat diet (HFD). Mice in the baseline group were sacrificed after four months of HFD feeding to establish advanced atherosclerosis. For the regression groups, mice were transitioned to a chow diet and treated with oral Lomitapide (MTP inhibitor) for an additional two weeks prior to sacrifice. Regression groups were further stratified into three subgroups based on intravenous treatments: (1) PBS, (2) VCAM1-targeting PROGRAMMED nanoparticles (Prog-VCAM1) encapsulating mutant PLPP3 mRNA (0.8 mg/kg, three injections), or (3) Prog-VCAM1 encapsulating functional PLPP3 mRNA (0.8 mg/kg, three injections). **(B)** Serum cholesterol levels are significantly reduced in the regression groups compared to the baseline group. n=6-8. Tukey’s post hoc test after one-way ANOVA. **(C)** Representative images of aortic root lesions stained with Oil Red O, CD68, and Sirius Red (bright-field and polarized light) from the baseline and regression groups. The regression groups comprise mice treated with Lomitapide and a chow diet for two weeks before sacrifice. Oil Red O staining quantifies plaque area, CD68 staining identifies macrophage content, and Sirius Red staining assesses collagen deposition. **(D)** The size of atherosclerotic plaques in the aortic root remains unchanged between the baseline and regression groups. n=6-8. Tukey’s post hoc test after one-way ANOVA. **(E)** CD68+ monocyte/macrophage accumulation within advanced atherosclerotic plaques is significantly reduced following treatment with Lomitapide and a chow diet, as quantified by the percentage of CD68+ area within the lesions. Intravenous administration of VCAM1-targeting PROGRAMMED nanoparticles encapsulating functional PLPP3 mRNA (Prog-VCAM1/PLPP3) further decreases CD68+ cell content compared to injections with PBS or VCAM1-targeting PROGRAMMED nanoparticles encapsulating mutant PLPP3 mRNA (Prog-VCAM1/PLPP3-Mut). This demonstrates significant therapeutic efficacy in mitigating inflammation within advanced lesions beyond the effects of aggressive cholesterol reduction, achieved through the delivery of functional PLPP3 mRNA by VCAM1-targeting PROGRAMMED nanoparticles. n=6-8. Tukey’s post hoc test after one-way ANOVA. **(F)** Collagen accumulation within atherosclerotic plaques increases following treatment with Lomitapide and a chow diet. Intravenous administration of VCAM1-targeting PROGRAMMED nanoparticles encapsulating functional PLPP3 mRNA (Prog-VCAM1/PLPP3) further enhances collagen deposition compared to treatment with PBS or VCAM1-targeting PROGRAMMED nanoparticles encapsulating mutant PLPP3 mRNA (Prog-VCAM1/PLPP3-Mut). This demonstrates significant therapeutic efficacy in stabilizing advanced lesions beyond the effects of aggressive cholesterol reduction, achieved through the targeted delivery of functional PLPP3 mRNA by VCAM1-targeting PROGRAMMED nanoparticles. n=6-8. Tukey’s post hoc test after one-way ANOVA. <u>All data are represented as mean ± SEM. *P < 0.05, **P < 0.01, ***P < 0.001.</u>

In the regression group, mice administered with mutant PLPP3 mRNA-encapsulated, VCAM1-targeting PROGRAMMED NP showed no change in lesion size compared to the baseline group (Fig. 5C and 5D). This group did exhibit decreased accumulation of CD68+ cells and increased collagen content (Fig. 5C and 5E). However, these lesion characteristics did not differ compared to the PBS-treated mice subjected to cholesterol lowering (Fig. 5C, 5D and 5E), showing VCAM1-targeting PROGRAMMED NP with nonfunctional mRNA had no therapeutic effect on lesion regression. In sharp contrast, mice in the regression group administered with functional PLPP3 mRNA-encapsulated, VCAM1-targeting PROGRAMMED NP exhibited markedly decreased accumulation of CD68+ cells (Fig. 5C and 5E) and increased collagen content (Fig. 5C and 5F) compared to PBS-treated mice and those subjected to mutant mRNA delivered by VCAM1-targeting PROGRAMMED NP. Weights (fig. S11) and plasma cholesterol levels (Fig. 5B) of mice remained consistent within all regression groups. These findings collectively demonstrate that functional PLPP3 mRNA-encapsulated VCAM1-targeting PROGRAMMED NP significantly enhances the regression of advanced atherosclerotic plaques, demonstrating additional therapeutic efficacy beyond the effects of aggressive cholesterol lowering. These results underscore the proof-of-concept of employing targeted mRNA nanomedicine to treat chronic vascular complications such as atherosclerotic diseases in patients who are currently undergoing control of systemic risk factors and lifestyle changes.

## Discussion

Vascular disease remains one of the most significant burdens on both human health and economic resources (*1, 44*). This challenge is further exacerbated by recent respiratory virus pandemics (*45, 46*) and the increasing prevalence of chronic vascular conditions influenced by lifestyle factors and demographic shifts toward aging populations (*47*). Our study highlights the potential of precision mRNA nanomedicine as a transformative approach for treating acute and chronic vascular diseases such as ARDS and atherosclerosis. Our approach of directly targeting human genetics-informed disease-causing mechanisms within inflamed endothelial cells shows considerable promise. We engineered and characterized the VCAM1-targeting PROGRAMMED NP, demonstrating its effectiveness in delivering functional mRNAs to intervene therapeutically in endothelial pathways informed by human genetics that drive the pathogenesis of ARDS and atherosclerosis.

Precision nanomedicine-assisted mRNA therapy has substantial potential for restoring cellular physiology and modulating or reversing disease processes by targeting critical and diverse molecular regulators of gene activity. Exploiting the inherent capability of mRNA to encode all known functional proteins, this strategy facilitates intervention across a spectrum of disease-causing biological pathways. Our investigation provides robust evidence for the effectiveness of mRNA delivery from genes with distinct molecular functions, exemplified by a transcription factor KLF2 for transcriptome regulation and a cell-surface glycoprotein PLPP3 for bioactive lipid hydrolysis. These targets present challenges for conventional pharmacological modulation. Transcription factors are compelling therapeutic targets due to their pivotal roles in orchestrating the expression of functionally interconnected genes (*48, 49*). Cell-state or tissue-specific dysregulation of transcription factors has been implicated in the pathogenesis of a wide array of diseases. Nevertheless, achieving precise delivery of therapeutic agents targeting transcription factors to specific cell types or tissues while mitigating off-target effects presents formidable technical challenges in contemporary drug development and delivery.

We have demonstrated here the transient restoration of physiological functions of the transcription factor KLF2 in diseased pulmonary microvasculature using targeted nanomedicine, resulting in a significant reduction of ARDS in mice. The loss of endothelial KLF2 not only contributes to ARDS but also exacerbates ventilator-induced lung injury (VILI) (*11, 12, 36*), which occurs due to mechanical ventilation commonly used in ARDS patients to support respiratory function (*50*). Therefore, the utilization of targeted KLF2 mRNA nanomedicine represents a promising adjunctive complementary therapeutic strategy for critically ill ICU patients with lung injury, encompassing conditions such as ARDS, VILI, sepsis, trauma, and others.

PLPP3 mRNA encodes phospholipid phosphatase 3, a cell-surface glycoprotein hydrolyzing phosphatidic acids. Endothelial loss of PLPP3, prompted by local disturbed blood flow and genetic predisposition, leads to regional vascular activation induced by extracellular lysophosphatidic acid (LPA) (*13–16*), a predominant bioactive lipid in circulation (*51, 52*). The VCAM1-targeting PROGRAMMED NP effectively restored PLPP3 expression in activated endothelial cells located in arterial regions prone to atherosclerosis. This intervention not only significantly attenuated the development of atherosclerotic lesions but also markedly promoted regression of advanced atherosclerotic plaques. Major adverse cardiovascular events (MACE) arising from plaque rupture and endothelial erosion of advanced arterial plaques persist as the primary life-threatening events globally, despite significant advancements in cholesterol-lowering therapies (*47*). Anti-inflammatory therapeutics, including IL-1β blockade, effectively reduce MACE, but are associated with side effects, notably an increased susceptibility to respiratory tract infections due to systemic inflammation inhibition (*53*). Our targeted mRNA nanomedicine approach provides spatiotemporal control of vascular inflammation, offering a vascular wall-targeted therapeutic strategy with potential to minimize side effects on systemic inhibition of inflammation.

Pipeline therapeutic targets associated with human genetic evidence of disease are twice as likely to result in approved drugs (*9, 10*). The proliferation of human genetics studies, including genome-wide association studies, has facilitated profound biological insights and therapeutic innovations, exemplified by the development of novel cholesterol-lowering agents such as PCSK9 inhibitors targeting hepatic tissues (*54, 55*). Our deliberate selection of KLF2 and PLPP3 mRNA for investigation stems from their known involvement in acute and chronic vascular inflammation, corroborated by robust evidence from molecular, cellular, and, more importantly, human genetics investigations (*11–14, 16*). Targeted nanomedicine serves as a pivotal conduit in translating genetically informed target identification and mechanistic *in vitro* and *in vivo* investigations into therapeutic development, especially given the tissue-specific nature of the majority of disease-causing mechanisms identified by GWAS (*56, 57*). For instance, the rs17114036 risk allele located on chromosome 1p32.2 exhibits one of the strongest associations with susceptibility to atherosclerosis by eliciting endothelial-specific suppression of PLPP3 transcription (*15, 16, 58*). This suggests that atherosclerotic patients with the rs17114036 risk allele could be prioritized for future vascular-targeted PLPP3 mRNA therapy as a form of personalized medicine.

The modularity of the PROGRAMMED NP formulation has the potential to spatiotemporally manipulate disease-causing mechanisms beyond inflamed endothelial cells. These platforms can display a diverse array of cell-targeting moieties and encapsulate various types of nucleotides. By utilizing DSPE-PEG, specific targeting moieties can be conjugated and presented on the corona of the PROGRAMMED NP, thereby facilitating the targeted delivery of therapeutic nucleotides to selected diseased cells and tissues.

In conclusion, we establish a proof-of-principle for vascular wall-targeted precision mRNA nanomedicine by devising inflamed-endothelium-targeting nanomaterials for the active delivery of functional mRNAs. This approach effectively targets human genetics-informed molecular mechanisms and treats acute and chronic vascular diseases *in vivo*, opening exciting possibilities in the field of vascular mRNA therapy.

## Acknowledgments

This work was supported by the National Institutes of Health (NIH) grants R35 HL161244 (Y.F.), R01 HL159558 (M.V.T.), and K99 HL166870 (Z.Z.); the American Heart Association (AHA) grants 23EIA1038679 (Y.F.) and 24POST1198682 (J.Z.); and the Gracias Family Foundation (Y.F., J.A.H., M.V.T.).

---

**Table S1.** Molar ratios of components in sixty nanoparticle formulations carrying functional KLF2 mRNA. The control nanoparticles encapsulate KLF2 mRNAs with mutations in all start codons (AUG), rendering them untranslatable.

| Round A screening |  |  |  |  |
| --- | --- | --- | --- | --- |
| Test No. | Molar ratios |  |  |  |
|  | G0-C14 | DOPE | Cholesterol | DSPE-PEG |
| A1 | 2 | 15 | 15 | 1 |
| A2 | 10 | 25 | 15 | 3 |
| A3 | 15 | 30 | 15 | 4 |
| A4 | 5 | 20 | 15 | 2 |
| A5 | 5 | 30 | 30 | 1 |
| A6 | 15 | 25 | 20 | 1 |
| A7 | 10 | 20 | 40 | 1 |
| A8 | 2 | 30 | 40 | 3 |
| A9 | 15 | 15 | 40 | 2 |
| A10 | 2 | 25 | 30 | 2 |
| A11 | 5 | 25 | 40 | 4 |
| A12 | 5 | 15 | 20 | 3 |
| A13 | 10 | 15 | 30 | 4 |
| A14 | 10 | 30 | 20 | 2 |
| A15 | 15 | 20 | 30 | 3 |
| A16 | 2 | 20 | 20 | 4 |
| A17 | 5 | 20 | 30 | 3 |
| A18 | 5 | 20 | 20 | 2 |
| A19 | 5 | 25 | 20 | 3 |
| Ctrl | 5 | 20 | 30 | 3 |

Round B screening
| Test No. | Molar ratios |  |  |  |
| --- | --- | --- | --- | --- |
|  | G0-C14 | DOPE | Cholesterol | DSPE-PEG |
| B1 | 2.5 | 20 | 15 | 2 |
| B2 | 5 | 30 | 15 | 4 |
| B3 | 5 | 25 | 15 | 3 |
| B4 | 2.5 | 15 | 30 | 4 |
| B5 | 2.5 | 30 | 20 | 2 |
| B6 | 5 | 20 | 15 | 1 |
| B7 | 2.5 | 25 | 15 | 4 |
| B8 | 2.5 | 25 | 20 | 1 |
| B9 | 2.5 | 20 | 30 | 3 |
| B10 | 2.5 | 30 | 15 | 3 |
| B11 | 5 | 30 | 30 | 1 |
| B12 | 2.5 | 15 | 15 | 1 |
| B13 | 5 | 25 | 30 | 2 |
| B14 | 5 | 15 | 15 | 2 |
| B15 | 5 | 20 | 20 | 4 |
| B16 | 5 | 15 | 20 | 3 |
| B17 | 10 | 15 | 30 | 2 |
| B18 | 5 | 20 | 35 | 3 |
| B19 | 5 | 20 | 20 | 3 |
| B20 | 5 | 20 | 25 | 2 |
| Ctrl | 5 | 20 | 30 | 3 |

Round C screening
| Test No. | Molar ratios |  |  |  |
| --- | --- | --- | --- | --- |
|  | G0-C14 | DOPE | Cholesterol | DSPE-PEG |
| C1 | 2.5 | 25 | 15 | 1 |
| C2 | 2.5 | 25 | 15 | 2 |
| C3 | 2.5 | 25 | 15 | 3 |
| C4 | 10 | 15 | 20 | 3 |
| C5 | 10 | 15 | 20 | 4 |
| C6 | 10 | 15 | 25 | 2 |
| C7 | 10 | 15 | 25 | 3 |
| C8 | 10 | 15 | 25 | 4 |
| C9 | 5 | 15 | 25 | 1 |
| C10 | 5 | 15 | 15 | 3 |
| C11 | 5 | 15 | 15 | 4 |
| C12 | 5 | 25 | 20 | 1 |
| C13 | 5 | 25 | 20 | 2 |
| C14 | 5 | 25 | 20 | 4 |
| C15 | 5 | 25 | 30 | 4 |
| C16 | 5 | 20 | 15 | 3 |
| C17 | 5 | 20 | 15 | 4 |
| C18 | 5 | 25 | 25 | 3 |
| C19 | 5 | 25 | 30 | 5 |
| C20 | 5 | 25 | 25 | 4 |
| C21 | 5 | 25 | 25 | 2 |
| Ctrl | 5 | 20 | 30 | 3 |

**fig. S1.**
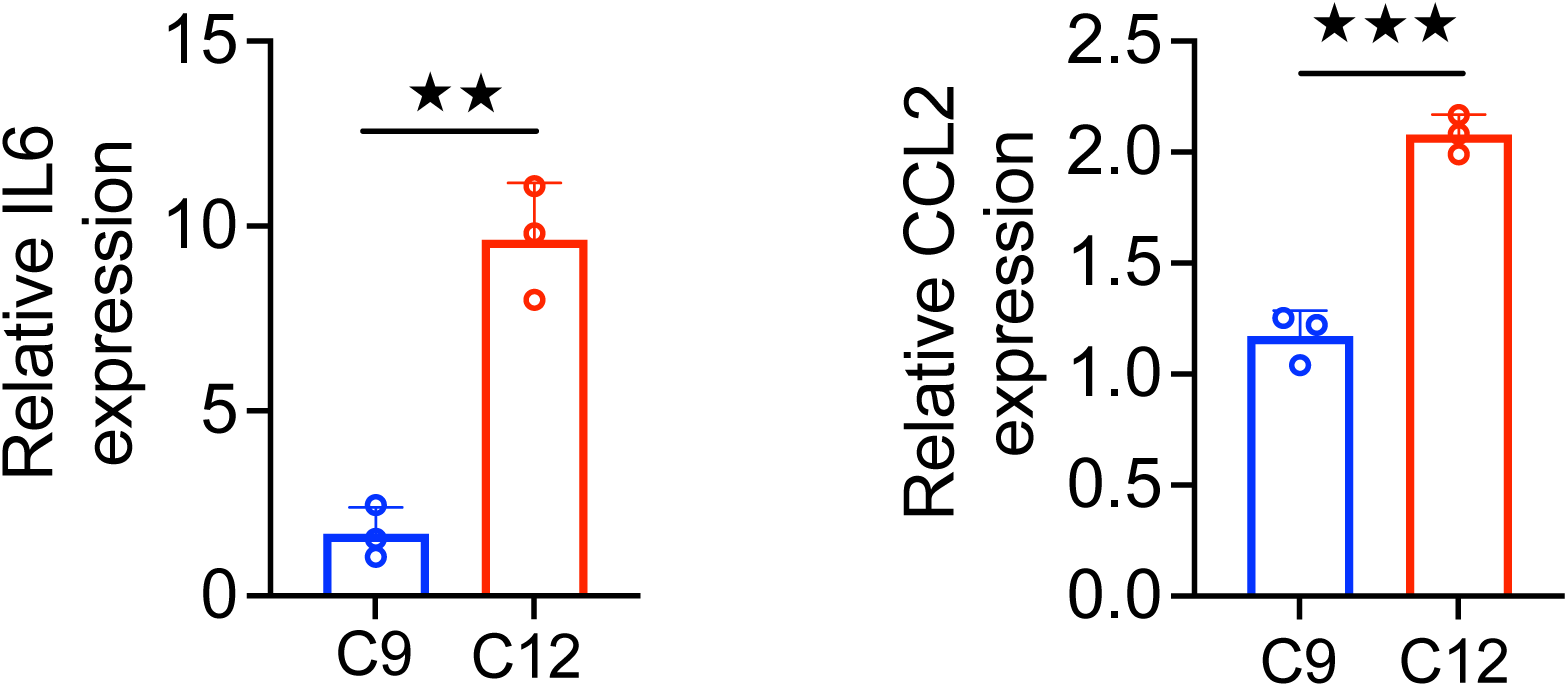
Expression of inflammatory biomarkers CCL2 and IL-6 in human pulmonary microvascular endothelial cells, quantified by real-time PCR, was elevated in cells treated with C12 nanoparticles compared to those treated with C9 nanoparticles (200 ng mRNA per well in a 24-well plate, 6 hrs). n=3. Student’s t-test.

**fig. S2.**
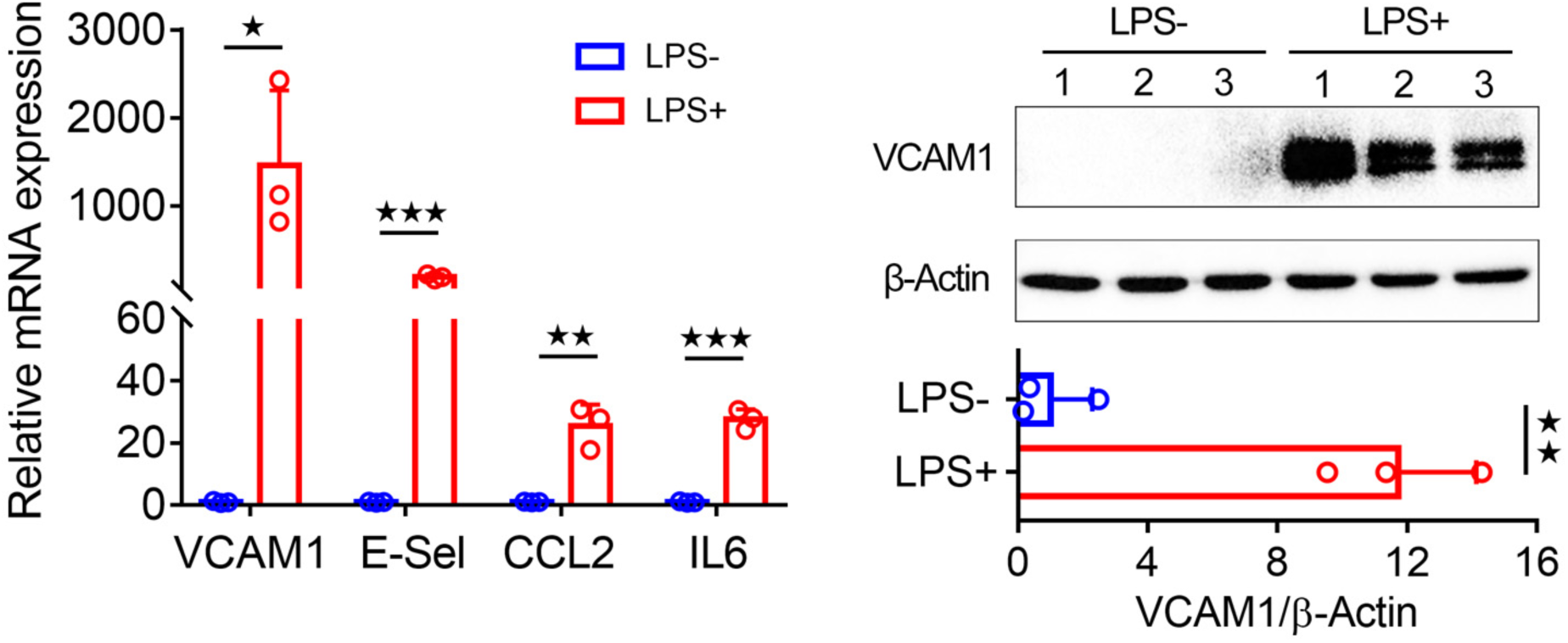
Lipopolysaccharides (LPS, 200 ng/mL, 3 hours) significantly induced inflammation in human pulmonary microvascular endothelial cells, including the significant up-regulation of vascular cell adhesion molecule 1 (VCAM-1). n=3. Student’s t-test.

**fig. S3.**
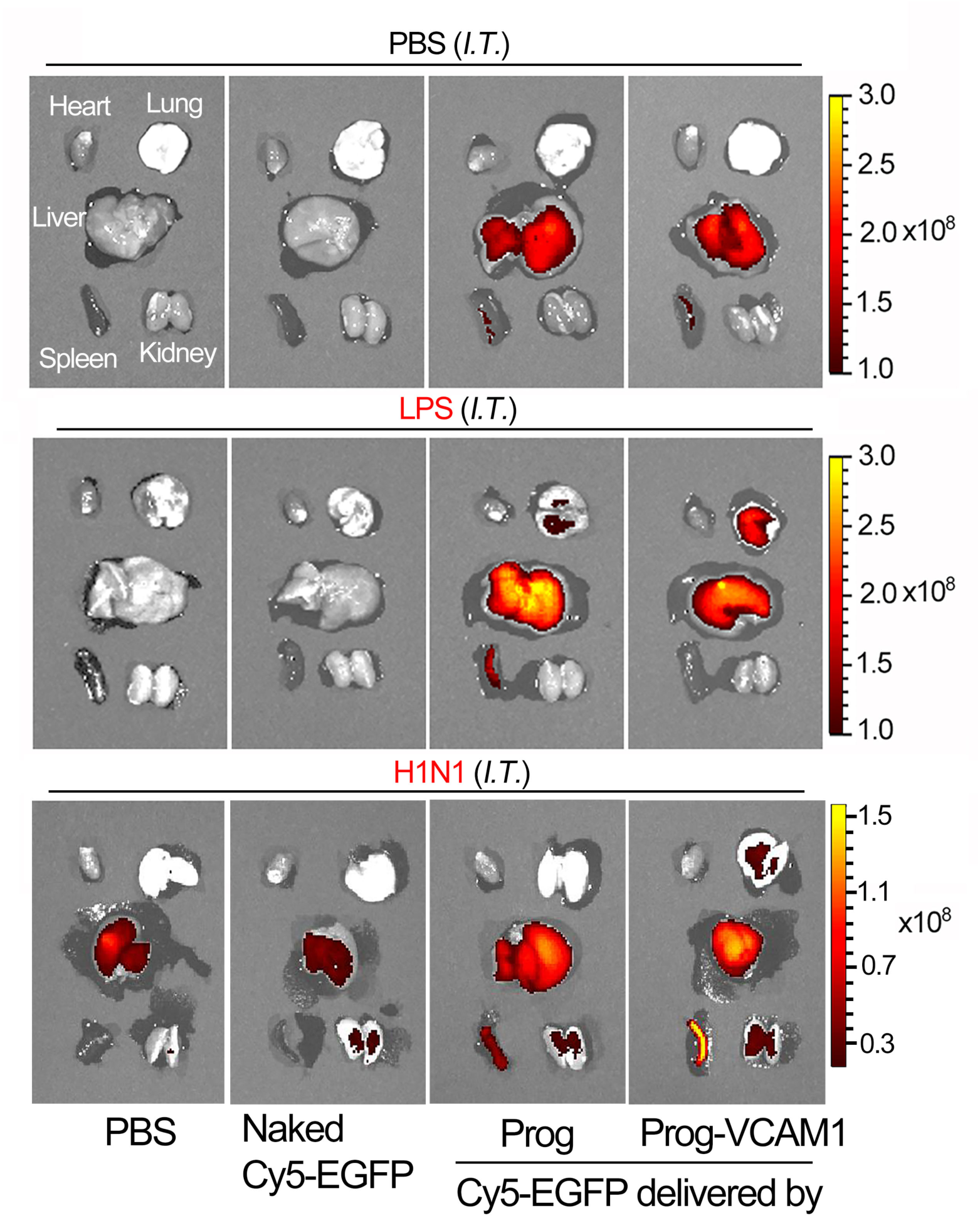
Representative biodistribution images of Cy5-labeled mRNA intravenously delivered in naked form or by non-targeting (Prog) or VCAM1-targeting PROGRAMMED nanoparticle (Prog-VCAM1) in mice subjected to intratracheal inoculation with PBS, LPS, or H1N1 virus. C57BL/6J mice were intratracheally instilled with PBS, LPS (50 uL, 1 mg/mL), or influenza A virus (50 uL, 200 PFU/mouse). One day later, they received intravenous injections of PBS, naked Cy5-labeled eGFP mRNA, or Cy5-labeled eGFP mRNA delivered by Prog or Prog-VCAM1. Major organs were excised and collected one day after NP injections, and Cy5 fluorescence imaging of organs was conducted using an IVIS 200 imaging system. Prog-VCAM1 enhanced mRNA delivery to ARDS lungs induced by LPS or H1N1 virus.

**fig. S4.**
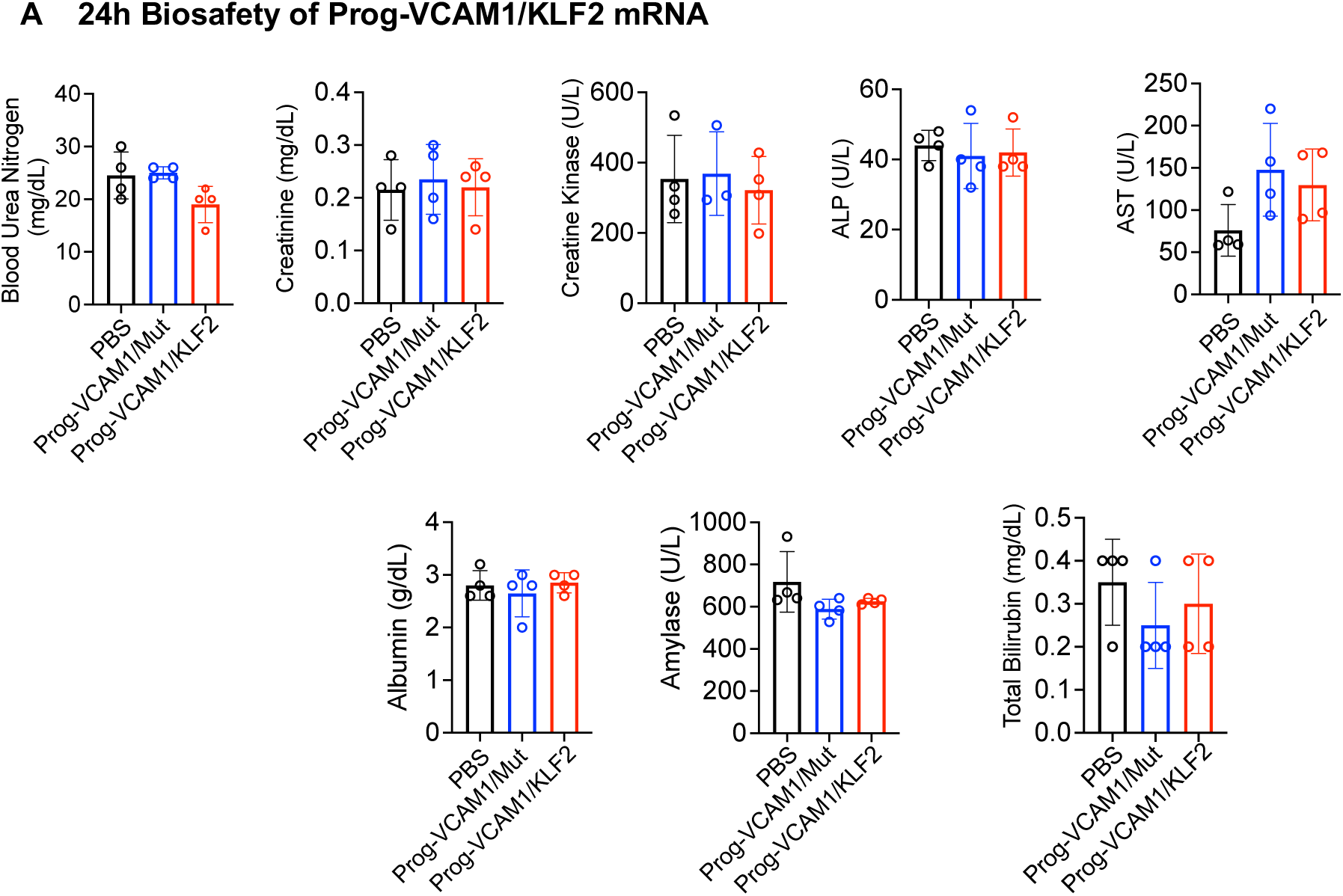
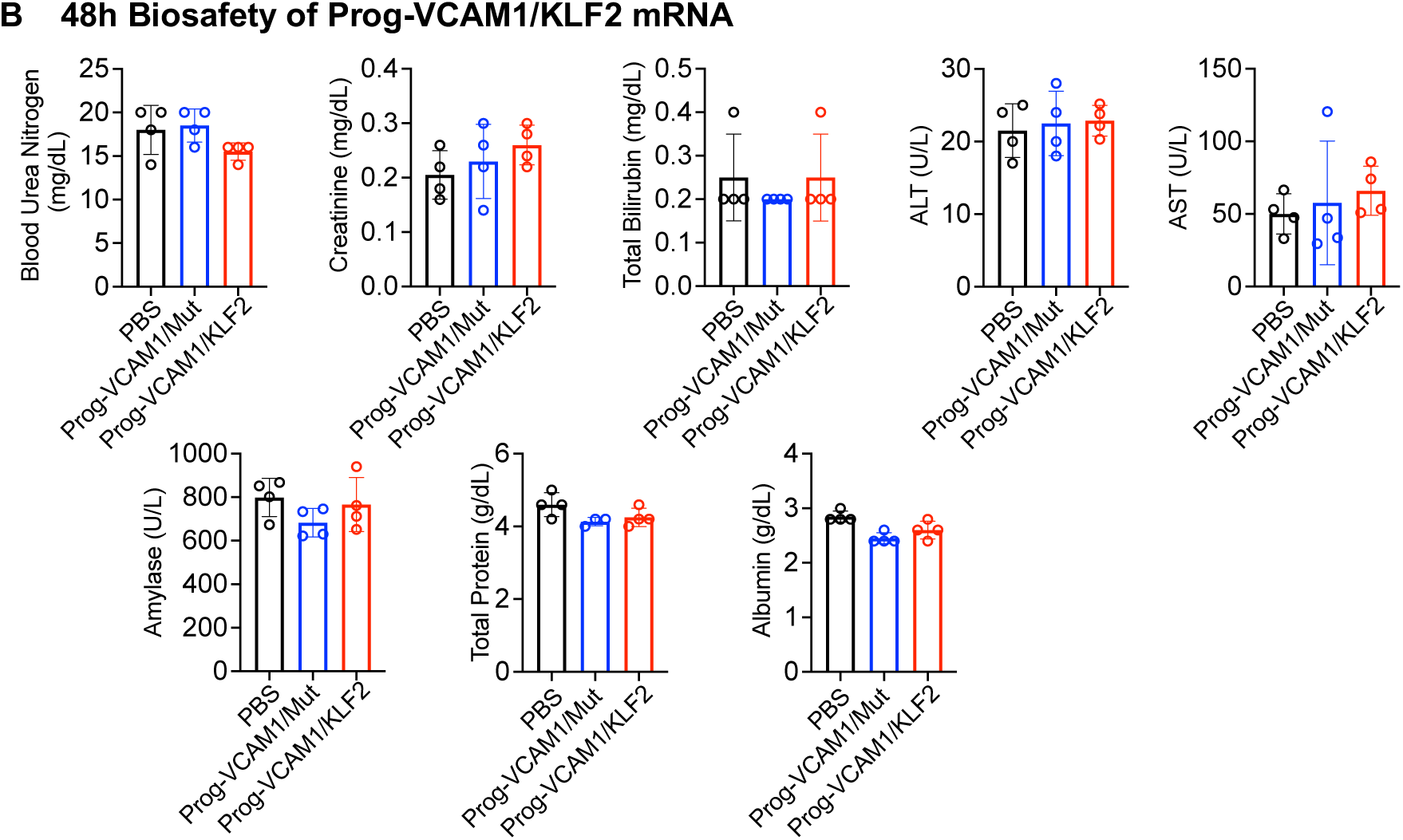
Assessment of physiological functions of liver, kidney, and pancreas following administration of VCAM1-targeting PROGRAMMED nanoparticle (Prog-VCAM1). Adult male C57BL/6J mice were intravenously injected with Prog-VCAM1 encapsulating mutant or functional KLF2 mRNA. Blood samples were collected at 24 and 48 hours post-injection to measure the activities of Alanine Aminotransferase (ALT), Aspartate Aminotransferase (AST), and Amylase, along with the concentrations of Blood Urea Nitrogen (BUN), Creatinine, Bilirubin, Albumin, and total protein. No significant changes in the function of the liver (ALT, AST, Bilirubin, Albumin, and total protein), kidney (BUN, Creatinine, Albumin, and total protein), and pancreas (Amylase) were detected **(A)** 24 and **(B)** 48 hours following administration of Prog-VCAM1. n=4. Student’s t-test.

**fig. S5.**
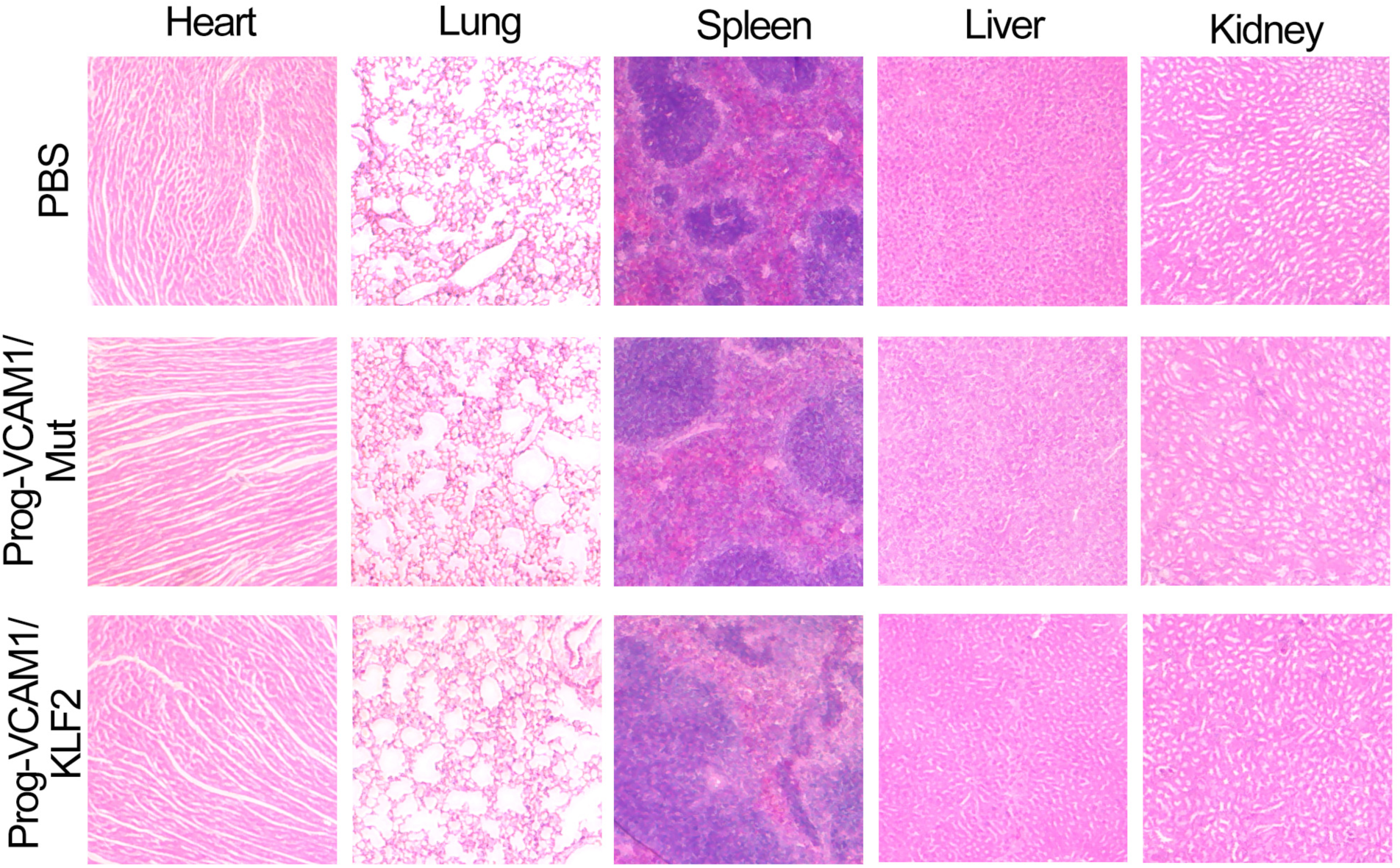
Histological sections of the heart, lung, spleen, liver, and kidney in C57BL/6J mice 48 hours following administration of PBS or VCAM1-targeting PROGRAMMED nanoparticle (Prog-VCAM1).

**fig. S6.**
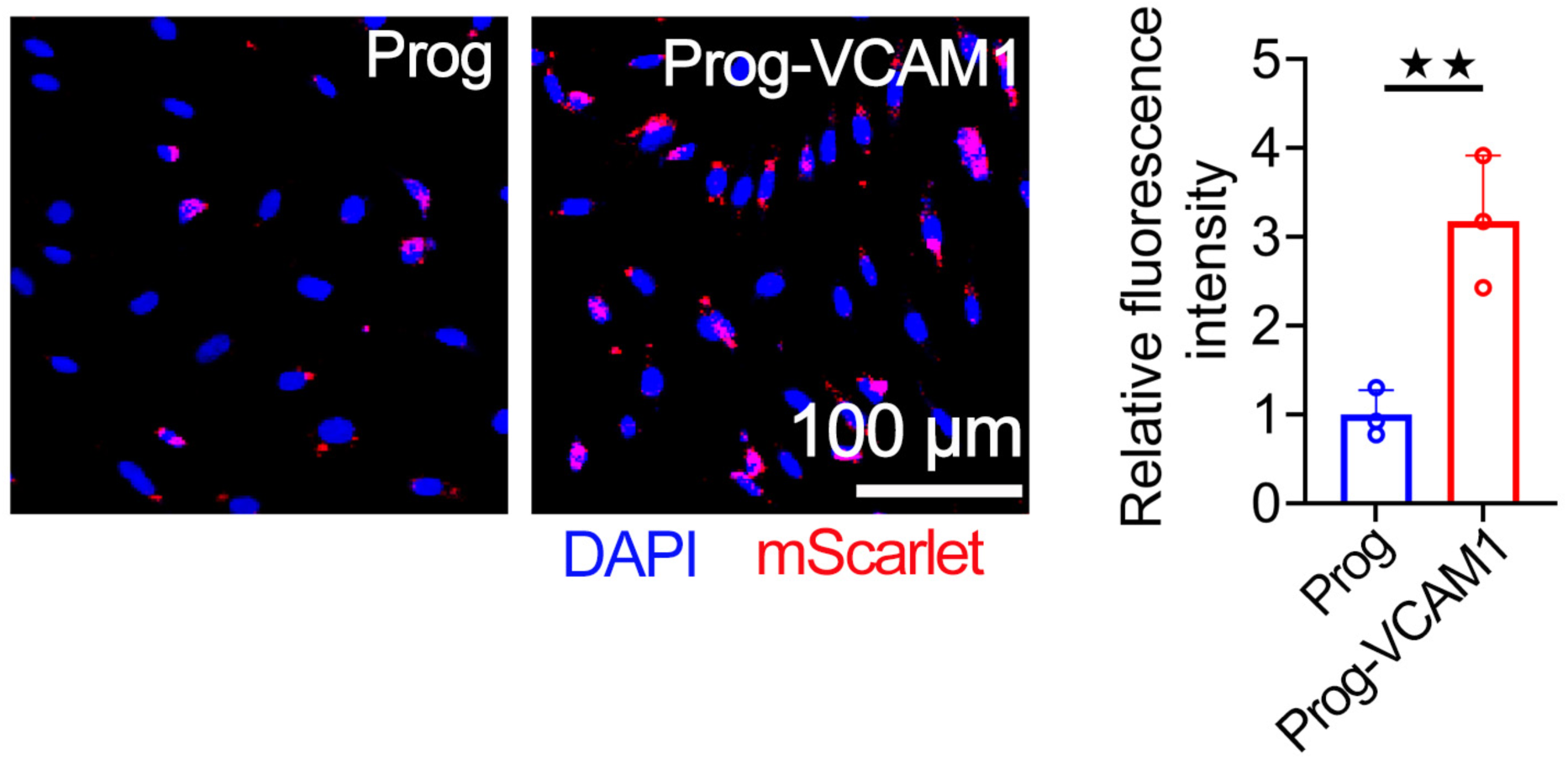
VCAM1-targeting PROGRAMMED nanoparticle (Prog-VCAM1) enhanced delivery of functional mScarlet mRNA to inflamed human aortic endothelial cells (HAECs). Inflammation in HAECs was induced by treatment with lipopolysaccharide (LPS, 200 ng/mL, 3 hours). Subsequently, HAECs were treated with either non-targeting Prog or targeting Prog-VCAM1 encapsulating m1Ψ mScarlet mRNA (200 ng mRNA per well in a 24-well plate). Fluorescence from mScarlet protein expression was quantified 6 hours after nanoparticle treatment. n=3. Student’s t-test.

**fig. S7.**
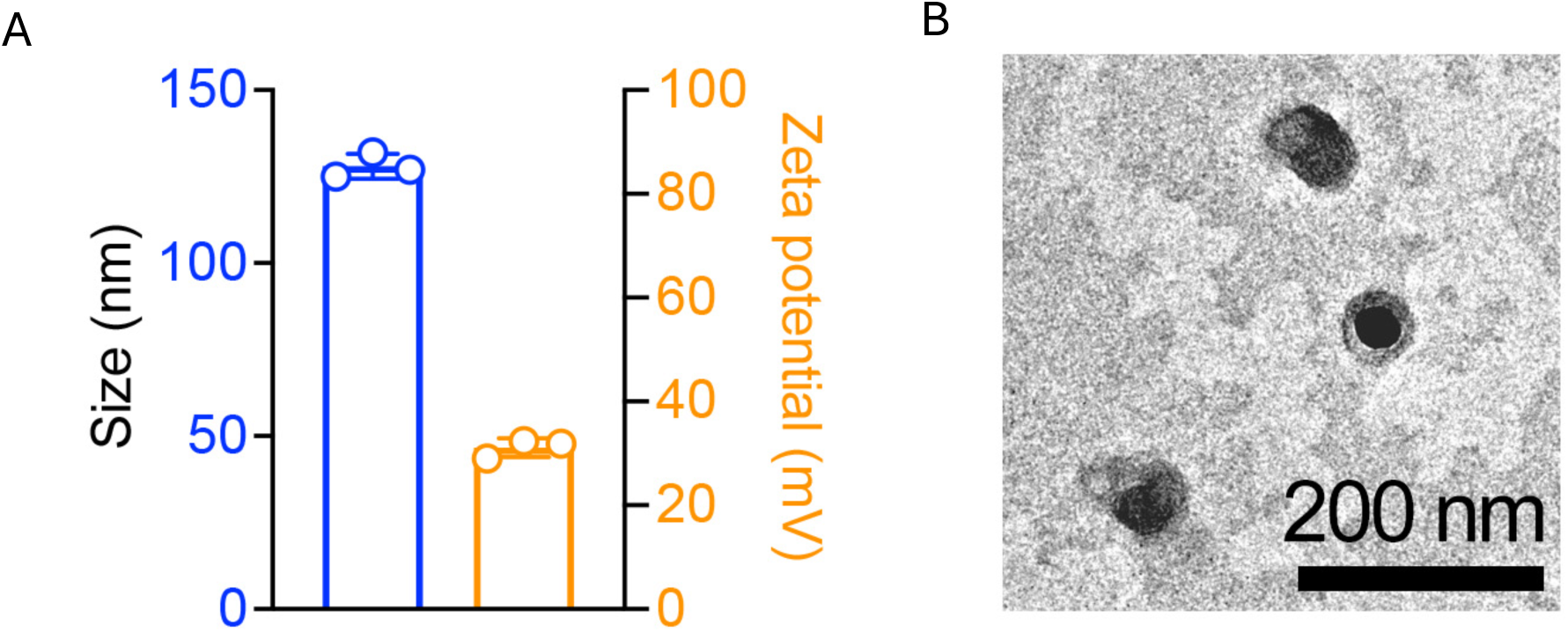
Hydrodynamic diameter size, Zeta potential (A) and negatively stained TEM image of PLPP3 mRNA-encapsulated VCAM1-targeting PROGRAMMED nanoparticle (Prog-VCAM1) (B). n=3. Student’s t-test.

**fig. S8.**
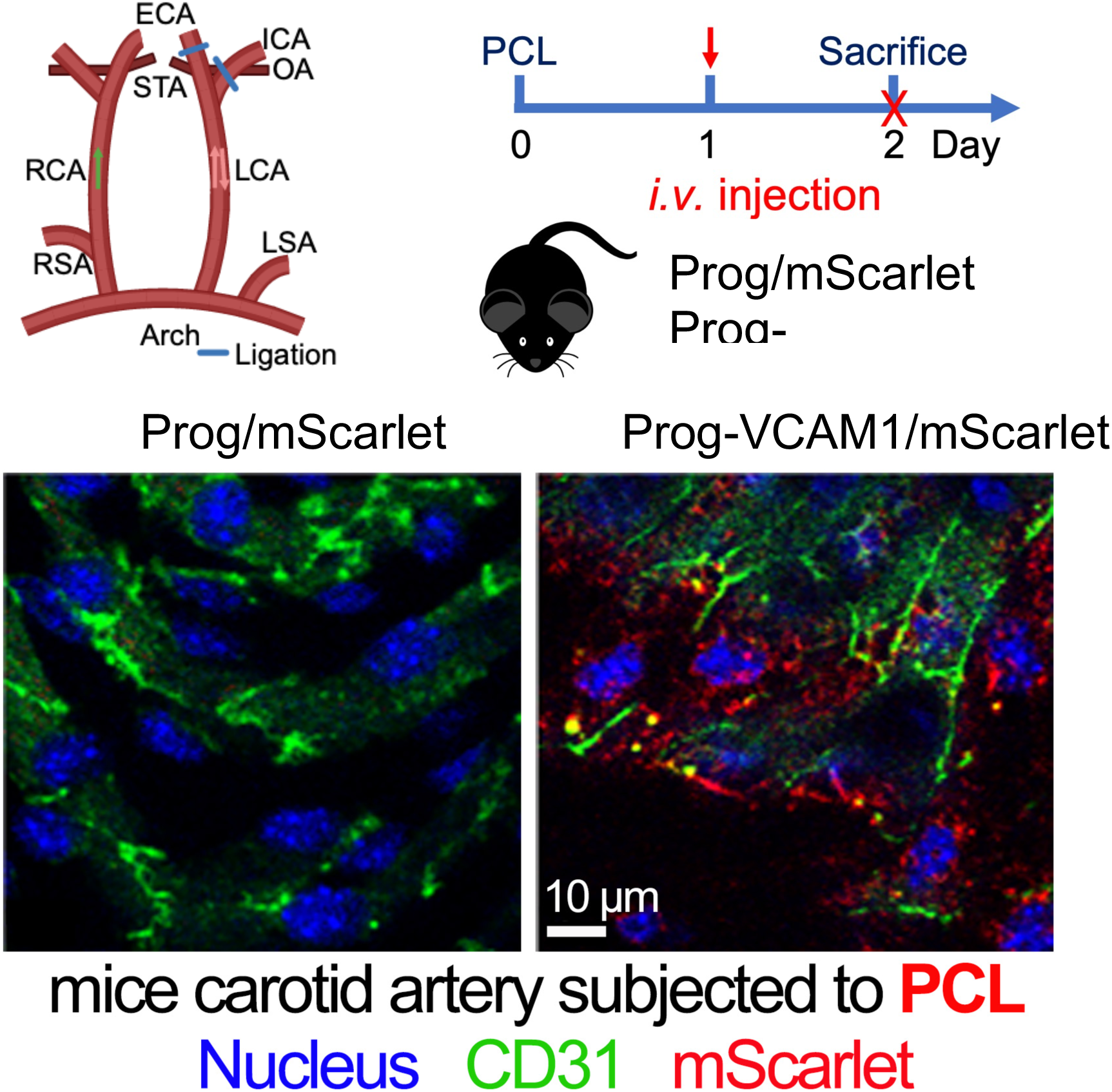
VCAM1-targeting PROGRAMMED nanoparticle (Prog-VCAM1), but not the non-targeting PROGRAMMED nanoparticle (Prog), effectively delivered functional mScarlet mRNA to inflamed endothelial cells in the ligated carotid artery of *Apoe*^−/−^ mice. Partial carotid artery ligation was performed in the left carotid artery (LCA) of high-fat-fed *Apoe*^−/−^ mice to induce acute disturbed flow and increase endothelial VCAM-1 expression. One day after ligation, m1Ψ mScarlet mRNA-encapsulated VCAM1-targeting or non-targeting Prog were intravenously administered. *En face* immunofluorescence images were obtained from mice sacrificed one day after NP injection.

**fig. S9.**
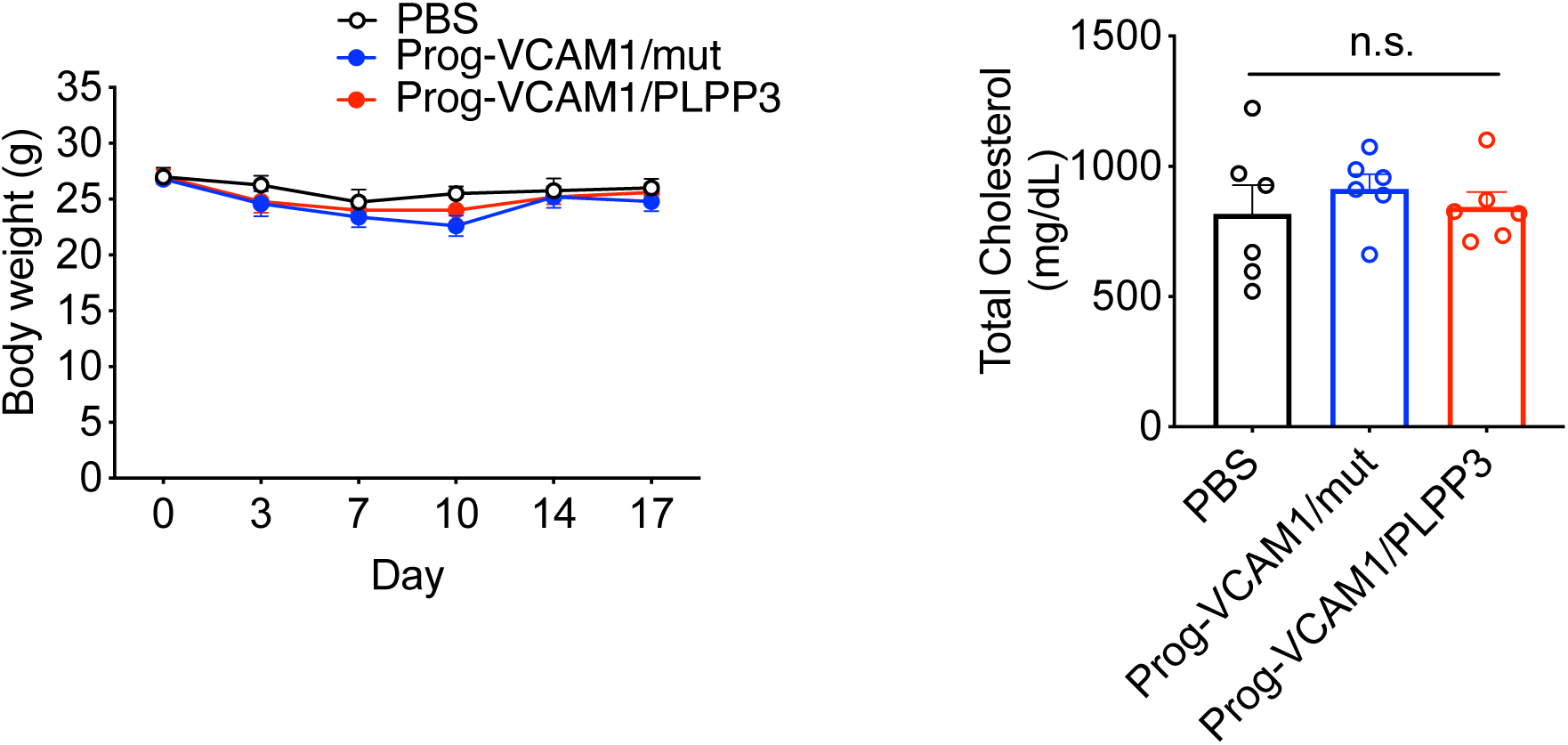
Body weights and plasma cholesterol levels were unaffected by injections of mRNA-encapsulated VCAM1-targeting PROGRAMMED nanoparticle (Prog-VCAM1) in *Apoe*^−/−^ mice subjected to carotid partial ligation. The study design is illustrated in Fig. 4E. n=5-6. Tukey’s post hoc test after one-way ANOVA.

**fig. S10.**
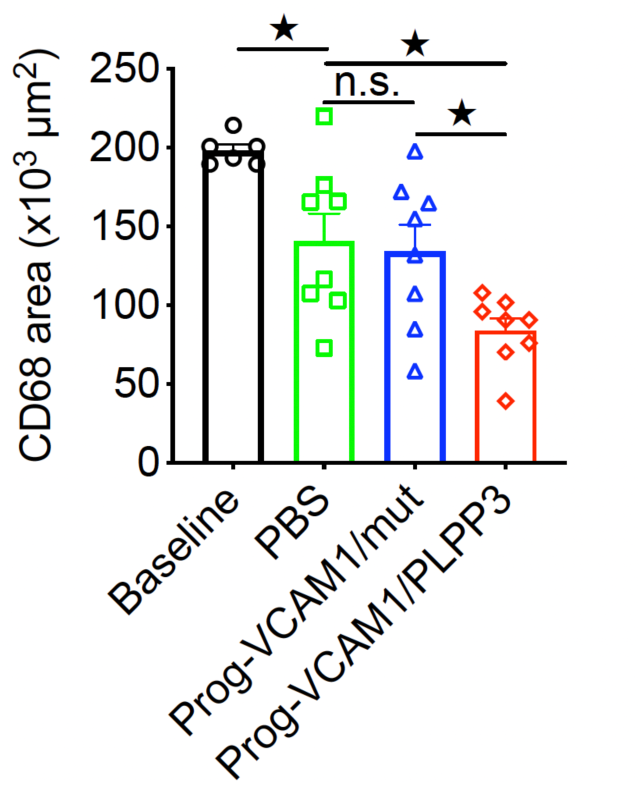
CD68+ monocyte/macrophage accumulation within advanced atherosclerotic plaques is significantly reduced following treatment with Lomitapide and a chow diet, as quantified by the total CD68+ area within the lesions. Intravenous administration of VCAM1-targeting PROGRAMMED nanoparticles encapsulating functional PLPP3 mRNA (Prog-VCAM1/PLPP3) further reduces CD68+ plaque area compared to injections with PBS or VCAM1-targeting PROGRAMMED nanoparticles encapsulating mutant PLPP3 mRNA (Prog-VCAM1/PLPP3-Mut). These results demonstrate significant therapeutic efficacy in mitigating inflammation within advanced lesions, beyond the effects of aggressive cholesterol reduction, achieved through the targeted delivery of functional PLPP3 mRNA by VCAM1-targeting PROGRAMMED nanoparticles. n=6–8. Tukey’s post hoc test after one-way ANOVA.

**fig. S11.**
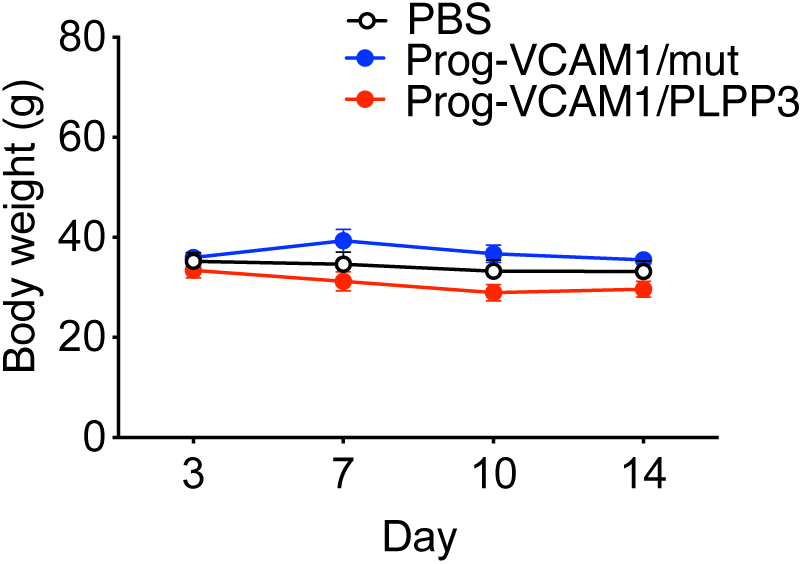
Body weights were unaffected by injections of mRNA-encapsulated VCAM1-targeting PROGRAMMED nanoparticle (Prog-VCAM1) in hypercholesteremic mice subjected to Lomitapide and chow diet treatment. The study design is illustrated in Fig. 5A. n=6-8. Tukey’s post hoc test after one-way ANOVA. <u>All data are represented as mean ± SEM. *P < 0.05, **P < 0.01, ***P < 0.001.</u>

## Supplementary Materials and Methods

### Materials and Methods

All studies were conducted in accordance with protocols approved by the Institutional Biosafety Committee (IBC) and the Institutional Animal Care and Use Committee (IACUC) of the University of Chicago. All experimental animals were assigned unique identifiers to blind experimenters to the treatments. A block randomization method was used to assign experimental animals to groups on a rolling basis, ensuring adequate sample sizes for each experimental condition.

#### Materials for Nanoparticles

VCAM1-targeting peptide (VHPKQHRGC) was purchased from GenScript (NJ, USA). 1,2-Epoxytetradecane and PAMAM dendrimer generation 0, Cholesterol, EDTA•2Na, and Tris(2-carboxyethyl) phosphine hydrochloride (TCEP) were purchased from Sigma-Aldrich (St. Louis, USA). DSPE-PEG-MAL (Mw=2000) and DSPE-mPEG (Mw=2000) were obtained from Nanosoft Polymers (NC, USA). 18:1 (Δ9-Cis) PE (DOPE) was purchased from Avanti Polar Lipids (AL, USA). Cy5-EGFP mRNA was obtained from APExBIO (TX, USA), and N1-methyl-pseudouridine (N1M) modified UTP (N1M-UTP) were purchased from TriLink BioTechnologies (CA, USA). Slide-A-Lyzer™ Dialysis Cassettes (20K MWCO) were purchased from Thermo Fisher (MA, USA).

#### *In vitro* transcription (IVT) of KLF2 and PLPP3 mRNA

The vector carrying the open reading frame (ORF) of human KLF2 with an HA tag was purchased from Addgene. The template for IVT of human KLF2 or PLPP3 mRNA, regulated by a T7 promoter, was amplified by PCR. IVT was conducted using a MEGAscript T7 kit (Ambion) containing 0.2 µg of the template, 7.5 mM ATP, 7.5 mM CTP, 7.5 mM N1M-UTP, 1.5 mM GTP, and 6.0 mM 3’-O-Me-m7G(5’)ppp(5’)G (ARCA). The reaction mixture was incubated at 37°C for 90 minutes, followed by a Turbo DNase digestion for 15 minutes. Poly(A) tails were then added to the mRNA using a Poly(A) kit (Ambion). The mRNA product was purified using a MEGAclear kit (Ambion) and resuspended in RNase-free water. Mutant, non-translatable KLF2 or PLPP3 mRNA were generated by IVT of the tramples in which all the translational start sites (AUG codon) were mutated. m1Ψ-modified mRNAs were generated by incorporating N1-methylpseudouridine (m1Ψ) in place of uridine during *in vitro* transcription. This RNA modification has been shown to enhance protein translation efficiency and attenuate innate immune activation.

#### Synthesis of Cationic Lipid G0-C14

G0-C14, a cationic lipid, was synthesized by reacting PAMAM G0 dendrimer with 1,2-epoxytetradecane according to established protocols (refs 1-2). 1,2-epoxytetradecane was added to the PAMAM G0 dendrimer at a mass ratio of 6:1. The reaction proceeded under vigorous stirring for 2 days. The resulting product was purified by silica chromatography using a gradient elution from CH₂Cl₂ to a CH₂Cl₂:MeOH:NH₄OH mixture (75:22:3).

#### Synthesis of DSPE-PEG-VHPKQHR

To synthesize DSPE-PEG-VHPKQHR, DSPE-PEG-MAL (100 mg) was dissolved in 3 mL of deionized (DI) water containing 40% ethanol. VCAM1-targeting peptide VHPKQHRGC (53 mg, dissolved in DI water), along with TCEP (20 mg, dissolved in DI water) and EDTA•2Na (3 mM, dissolved in DI water), were added to the solution. The reaction was allowed to proceed for 24 hours under a nitrogen (N₂) atmosphere. Following the reaction, the product was purified by extensive dialysis (MWCO = 1000) against DI water and subsequently lyophilized to obtain DSPE-PEG-VHPKQHR.

#### PROGRAMMED nanoparticle formulation

Lipids containing G0-C14, DOPE, Cholesterol, and DSPE-PEG were dissolved in ethanol at specified molar ratios according to the designed formulation table to assemble non-targeting LNPs. The VCAM1-targeting LNPs were formulated identically, except DSPE-PEG was replaced with DSPE-PEG-VCAM1. The lipids were mixed with nuclease-free DI water containing mRNA at a volume ratio of 3:1 (aqueous:ethanol = 3:1, v/v). The nitrogen to phosphate ratio (N:P) on the cationic lipid was set to 8.5 for each formulation. The formed LNPs were then dialyzed with sterilized PBS for 2 hours to remove ethanol using a Slide-A-Lyzer dialysis cassette with a MWCO of 20K.

#### Nanoparticle characterization

Transmission electron microscope (TEM) images were obtained using an FEI Spirit LaB6 Routine Electron Microscope operating at 300 kV, located in the Advanced Electron Microscopy Facility at The University of Chicago. NMR spectra were recorded using a 400 MHz NMR instrument (Bruker, Germany). The hydrodynamic size and zeta potential were measured using a Zetasizer Nano ZS90 (Malvern, UK).

#### Cell culture

Human macrovascular endothelial cells (HMVECs) and human aortic endothelial cells (HAECs) were obtained from Lonza (NJ, USA) and cultured in EGM2 medium containing 2% FBS at 37°C with 5% CO2. Cells at passages 5-8 were used for the experiments.

#### Influenza A virus

The following reagent was obtained through BEI Resources, NIAID, NIH: Influenza A Virus, A/Puerto Rico/8/1934 (H1N1), NR-348.The virus was stored at -80°C.

#### Quantitative real-time PCR (qPCR)

For cultured cells, total RNA was extracted using the GenElute Mammalian Total RNA Miniprep Kit (Sigma). For RNA extraction from mouse tissues, the tissues were first homogenized with a motorized tissue grinder (Fisherbrand, Fisher Scientific) and then processed using the chloroform-isopropanol method. A total of 0.2 µg of mRNA was reverse-transcribed into cDNA using the High-Capacity cDNA Reverse Transcription Kit (Life Technologies). The cDNA was amplified using a LightCycler 480 II (Roche) system with SYBR Green as the probe. Absolute quantification of the gene of interest was normalized to the geometric mean of β-actin, GAPDH, and ubiquitin.

#### *In vitro* optimization of PROGRAMMED nanoparticles

HMVECs were seeded in a 24-well plate overnight to reach 80-100% confluence before the experiments. A library of sixty PROGRAMMED nanoparticles with different molar ratios of components was formulated and screened for their capacity to effectively deliver functional mRNA to endothelial cells (Table S1). The control nanoparticle encapsulated KLF2 mRNAs with mutations in all start codons (AUG), rendering them untranslatable. Cells were transfected with nanoparticles at a KLF2 mRNA concentration of 200 ng for 24 hours. The optimized C9 PROGRAMMED NP formulation, determined by the up-regulation of RAPGEF3 transcripts compared to cells treated with the control nanoparticle, was identified from the library through three rounds of mRNA transfection screens and selected for subsequent studies.

#### Cellular uptake of the PROGRAMMED nanoparticles

HMVECs were seeded in a 24-well plate overnight to reach 80-100% confluence. Before mRNA NP incubation, HMVECs were pre-treated with one of the following inhibitors for 1 hour: chlorpromazine (20 µM), wortmannin (20 nM), cytochalasin D (15 µM), or genistein (500 µM). The inhibitors were then removed, and HMVECs were transfected with mRNA NPs containing 0.2 µg of Cy5-EGFP mRNA for 1 hour. The cells were washed and resuspended in PBS for flow cytometry analysis. For microscopy imaging, the cells were washed and fixed after nanoparticle incubation for 1 hour and stained with Hoechst 33342 (2 µg/mL in PBS) for nuclear identification. Imaging was performed using a confocal laser scanning microscope (CLSM, Leica SP5 2-Photon, Germany).

#### VCAM1-targeting PROGRAMMED NP for functional mRNA delivery to inflamed endothelial cells *in vitro*

HMVECs were seeded in a 24-well plate overnight to reach 80-100% confluence before experiments. Endothelial inflammation was induced by treating cells with LPS (200 ng/mL) for 3 hours at 37 °C, which was verified by a marked increase in E-selectin, CCL2, IL6, and VCAM-1 expression, as analyzed by qPCR. PBS- or LPS-treated HMVECs were then incubated with either non-functionalized C9 PROGRAMMED NP or VCAM1-targeting PROGRAMMED NP, each encapsulating KLF2 mRNA (0.2 µg) or control KLF2 mutant mRNA. After 30 minutes of incubation, the culture medium was removed and replaced with 500 µL of fresh culture medium. Cells were harvested 6 hours later, and the expression of the KLF2 downstream gene *NOS3* was analyzed by qPCR. For imaging and quantification of functional mRNA delivery, HMVECs were seeded in µ-slide 8-well glass-bottom dishes (Ibidi, 80827) and incubated for 24 hours, followed by LPS treatment (200 ng/mL) for 3 hours at 37 °C to induce inflammation. The cells were then treated with non-functionalized C9 PROGRAMMED NP or VCAM1-targeting NP PROGRAMMED encapsulating 0.2 µg mScarlet mRNA. After 30 minutes of nanoparticle incubation, the culture medium was removed, and cells were washed with PBS. To fix the cells, 200 µL of paraformaldehyde (4% in PBS) was added to each well for 15 minutes, followed by nuclear staining with Hoechst 33342 (2 µg/mL in PBS) for 10 minutes. Fluorescence images were captured using a 40x oil immersion objective on a confocal laser scanning microscope (CLSM)

#### Plasma terminal half-life of NP-delivered mRNA *in vivo*

Adult male C57BL/6J mice were intravenously injected via the tail vein, with Cy5-EGFP mRNA (0.8 mg/kg body weight) in either naked form or encapsulated in non-functionalized C9 PROGRAMMED NP or VCAM1-targeting PROGRAMMED NP. Blood samples (∼50-100 μL) were collected at predetermined intervals (0, 30, 60, 120, and 240 minutes) post-injection via submandibular bleeding. Plasma was isolated by centrifuging the blood at 5000 rpm for 10 minutes. The fluorescence intensity of Cy5-EGFP mRNA in plasma was measured at emission and excitation wavelengths of 670 nm and 640 nm, respectively, after diluting 15 μL of supernatant with 45 μL PBS. Pharmacokinetics were determined by normalizing the fluorescence intensity of each time point to that of the initial (0 min) time point.

#### Biodistribution of NP-delivered functional mRNA in mouse models of ARDS *in vivo*

To induce lung inflammation, adult male C57BL/6J mice were treated intratracheally with LPS (1 mg/kg in PBS) for 6 hours. The mice were then intravenously injected with Cy5-EGFP mRNA (0.8 mg/kg body weight), either in naked form or encapsulated within non-functionalized C9 PROGRAMMED NP or VCAM1-targeting PROGRAMMED NP. Eighteen hours post-injection, the mice were euthanized, and the heart, lung, liver, spleen, and kidney were harvested for biodistribution analysis of the nanoparticles. Cy5 fluorescence imaging of organs was measured using an *in vivo* imaging system (IVIS 200, PerkinElmer). To evaluate the biodistribution of NP-delivered mRNA in H1N1 virus-infected mice, adult male C57BL/6J mice were treated intratracheally with influenza virus H1N1 (200 PFU per mouse in PBS) for 24 hours. The mice were then injected with Cy5-EGFP mRNA (0.8 mg/kg body weight), either in naked form or encapsulated within non-functionalized C9 PROGRAMMED NP or VCAM1-targeting PROGRAMMED. Twenty-four hours post-injection, the mice were euthanized, and the heart, lung, liver, spleen, and kidney were harvested for biodistribution analysis. Cy5 fluorescence imaging of organs was measured using an *in vivo* imaging system (IVIS 200, PerkinElmer).

#### Biosafety

The biosafety of VCAM1-targeting PROGRAMMED nanoparticles was assessed by analyzing key biomarkers of organ function and performing histological evaluations of major tissues. Adult male C57BL/6J mice (8–10 weeks old) were intravenously injected with VCAM1-targeting PROGRAMMED NPs encapsulating either mutant or functional KLF2 mRNA at a specified dose. Control mice received an equivalent volume of PBS. Blood samples were collected at 24 and 48 hours post-injection to evaluate serum biomarkers associated with liver, kidney, and pancreatic function, as well as overall protein balance using a biochemistry analyzer (Alfa Wassermann Diagnostic Technologies) according to the manufacturer’s instructions. Serum biochemical analyses were conducted to measure the activities of alanine aminotransferase (ALT), aspartate aminotransferase (AST), and amylase, along with the concentrations of blood urea nitrogen (BUN), creatinine, bilirubin, albumin, and total protein. Liver function biomarkers, including ALT, AST, and bilirubin, showed no significant differences between NP-treated and PBS-treated mice, indicating no hepatotoxicity. Kidney function was also unaffected, as evidenced by comparable levels of BUN and creatinine in both groups. Similarly, pancreatic function remained normal, with consistent amylase activity observed across treatment groups. Furthermore, the levels of albumin and total protein remained unchanged, confirming the preservation of liver and kidney function in NP-treated mice. To further investigate potential tissue damage, histological analyses of the heart, lungs, spleen, liver, and kidneys were performed 48 hours post-administration. Organs were harvested, fixed in 10% neutral-buffered formalin, embedded in paraffin, sectioned, and stained with hematoxylin and eosin (H&E). Microscopic examination revealed no structural abnormalities, inflammation, or other pathological changes in NP-treated mice compared to PBS controls.

#### *In vivo* delivery of functional mRNA to inflamed pulmonary microvascular endothelial cells is achieved using VCAM1-targeting PROGRAMMED nanoparticles

To induce lung inflammation, adult male C57BL/6J mice were treated intratracheally with LPS (1 mg/kg in PBS) for 6 hours. The mice were then intravenously injected via the tail vein, with either non-functionalized non-targeting PROGRAMMED NP or VCAM1-targeting PROGRAMMED NP encapsulating m1Ψ-modified mScarlet mRNA (1 mg/kg body weight). Eighteen hours post-injection, the mice were euthanized, and lungs were harvested and embedded in paraffin. Tissue sections were deparaffinized, rehydrated, and rinsed with distilled water. Antigen retrieval was performed by boiling sections in citrate buffer (pH 6) for 10 minutes, followed by treatment with 0.3% hydrogen peroxide to quench endogenous peroxidase activity, and permeabilization with 0.1% Triton X-100. After blocking for 1 hour, the slides were incubated overnight at 4 °C with Alexa Fluor 488-labeled CD31 antibody (R&D, FAB6874G) to label endothelial cells. After washing with PBS, the slides were mounted with fluorescence medium containing DAPI (Abcam) for nuclear counterstaining, and fluorescence images were acquired using a confocal laser scanning microscope (CLSM, Leica SP5 2-Photon, Germany). To assess mRNA delivery to inflamed pulmonary microvascular endothelial cells in H1N1-infected mouse lungs, adult male C57BL/6J mice were treated intratracheally with influenza virus H1N1 (200 PFU per mouse in 50 µL PBS) for 24 hours. The mice were then intravenously injected with either non-functionalized PROGRAMMED NP or VCAM1-targeting PROGRAMMED NP encapsulating mScarlet mRNA (1 mg/kg body weight). Four hours after nanoparticle injection, the mice were euthanized, and the lungs were harvested. Lung sections and subsequent immunohistochemical staining were conducted following the same procedure described above. Lung tissues were also collected for RNA isolation and quantitative real-time PCR (qRT-PCR) analysis.

#### Therapeutic effectiveness of KLF2 mRNA-encapsulated VCAM1-targeting PROGRAMMED NP in lessening ARDS in mice

To induce acute respiratory distress syndrome (ARDS), adult male C57BL/6J mice were treated intratracheally with influenza virus H1N1 (200 PFU in PBS) on day 0. The mice were then intravenously injected via the tail vein with VCAM1-targeting PROGRAMMED NP encapsulating either m1Ψ-modified KLF2 mRNA or mutant KLF2 control mRNA (0.8 mg/kg body weight) on days 1 and 4. On day 6, the mice were euthanized using ketamine (100 mg/kg) and xylazine (10 mg/kg).Bronchoalveolar lavage (BAL) was performed with 0.75 mL PBS containing 0.1 mM EDTA, and total cell counts in the bronchoalveolar lavage fluid (BALF) were determined using a hemocytometer. Total protein concentration in BALF was measured with a BCA protein assay kit, and pro-inflammatory cytokine levels in BALF and lung tissues were assessed using the LEGENDplex mouse cytokine release syndrome panel kit (BioLegend). The left lung lobe was harvested for RNA and protein analysis, while the right lobe was fixed in 4% paraformaldehyde (PFA) for histological examination. Lung injury severity was scored on paraffin-embedded lung sections using a histological scoring system (*1, 2*), graded as follows: 0: no damage; 1: mild; 2: moderate; 3: severe; and 4: very severe injury, with an increment of 0.5 if the lung injury fell between two integers.

#### VCAM1-targeting PROGRAMMED NP for functional PLPP3 mRNA delivery to inflamed endothelial cells *in vitro*

The endothelial tropism of non-targeting PROGRAMMED nanoparticles (NPs) was assessed by evaluating their capacity to deliver functional m1Ψ-modified PLPP3 mRNA to cultured human aortic endothelial cells (HAECs). HAECs were seeded in 24-well plates and cultured overnight to reach 80–100% confluence before experimentation. Cells were transfected with non-targeting PROGRAMMED NPs encapsulating either functional m1Ψ-modified PLPP3 mRNA (flag-tagged) or mutant PLPP3 mRNA (200 ng) for 6 hours. Following transfection, cells were harvested for quantitative real-time PCR (qRT-PCR) and Western blot analysis to assess mRNA expression and protein production. To evaluate the delivery efficiency of non-targeting and VCAM1-targeting PROGRAMMED NPs under inflammatory conditions, HAECs were pre-treated with lipopolysaccharide (LPS, 200 ng/mL) for 3 hours at 37 °C to induce inflammation. Inflamed HAECs were subsequently treated with either non-targeting PROGRAMMED NPs or VCAM1-targeting PROGRAMMED NPs encapsulating m1Ψ-modified mScarlet mRNA (200 ng) for 6 hours. Following treatment, cells were fixed, and fluorescence imaging was performed to assess mRNA delivery and the resultant protein expression.

#### *In vivo* delivery of functional PLPP3 mRNA to inflamed arterial endothelial cells using VCAM1-targeting PROGRAMMED NP

To induce acute disturbed flow, partial carotid artery ligation (PCL) was performed on the left carotid artery (LCA) of high-fat diet-fed male *Apoe^−/−^* mice, resulting in increased VCAM-1 expression and decreased PLPP3 expression in endothelial cells compared to the non-ligated right carotid artery (RCA) (*3, 4*). To evaluate the efficacy of VCAM1-targeted PROGRAMMED nanoparticles for delivering m1Ψ-modified PLPP3 mRNA to inflamed arterial endothelial cells *in vivo*, mice received one intravenous tail vein injection of PBS or m1Ψ-modified PLPP3 mRNA (0.8 mg/kg body weight) encapsulated within non-targeting or VCAM1-targeting PROGRAMMED NP one day post-ligation. Twenty-four hours post-injection, carotid artery tissues (ligated and non-ligated) were harvested, and endothelium-enriched intimal RNA was extracted using QIAzol lysis reagent (QIAGEN, MD, USA) following established protocols (*3, 4*). PLPP3 mRNA expression levels were quantified by real-time PCR. For protein-level assessment, male *Apoe^−/−^* mice subjected to PCL received a single intravenous injection of PBS, VCAM1-targeting PROGRAMMED NP encapsulating mutant PLPP3 mRNA, or functional PLPP3 mRNA tagged with a Flag (DYKDDDDK)-expressing sequence (2 mg/kg body weight) one day post-ligation*. En face* immunofluorescence staining of carotid arteries was performed 24 hours post-injection to detect Flag-tagged PLPP3 protein expression. Both ligated and contralateral carotid arteries were dissected and fixed in 4% paraformaldehyde for 15 minutes, then opened longitudinally to expose the endothelium. After permeabilization with 0.3% Triton X-100 for 30 minutes and blocking with 2% BSA for 1 hour at room temperature, the arteries were incubated overnight at 4°C with primary antibodies (Anti-Flag, #NBP1-06712, Novus; VE-Cadherin, #ab33168, Abcam). Following PBS washes, arteries were incubated with Alexa Fluor 594-labeled anti-rabbit and Alexa Fluor 647-labeled anti-rat secondary antibodies (Invitrogen) for 1 hour at room temperature. After additional PBS washes, nuclei were stained with DAPI (1:1000) for 10 minutes. The vessels were mounted on slides using anti-fade mounting medium (Abcam, UK) with the endothelium facing the cover glass. Fluorescence imaging was conducted using a 3i Marianas Spinning Disk Confocal Microscope, and images were analyzed with Slidebook software (Intelligent Imaging Innovations, CO, USA). Additionally, *en face* imaging of the ligated carotid artery was conducted in *Apoe^−/−^* mice one day after intravenous injection of m1Ψ-modified mScarlet RNA (2 mg mRNA/kg body weight) encapsulated within either non-targeting PROGRAMMED NP or VCAM1-targeting PROGRAMMED NP.

#### Therapeutic effectiveness of PLPP3 mRNA-encapsulated VCAM1-targeting PROGRAMMED NP in reducing atherosclerosis progression in mice

Adult male *Apoe^−/−^* mice were fed a high-fat diet (TD. 88137, Harlan-Envigo, IN, USA) and subjected to partial carotid artery ligation (PCL) to induce disturbed flow and vascular inflammation in the left carotid artery (LCA). Atherosclerosis in the ligated LCA was confirmed 14 days post-surgery. The ligated arteries were excised, fixed in 4% paraformaldehyde for 15 minutes, dehydrated overnight in 30% sucrose, and embedded in optimal cutting temperature (OCT, Tissue Tek) compound. Serial cross-sections (8 μm) were prepared and stained with Oil Red O to visualize atherosclerotic plaques.To assess the therapeutic potential of PLPP3 mRNA-encapsulated nanoparticles, mice were administered three intravenous injections (on days 1, 6, and 11 post-PCL) of either PBS, VCAM1-targeting PROGRAMMED nanoparticles encapsulating mutant control m1Ψ-modified PLPP3 mRNA, or VCAM1-targeting PROGRAMMED nanoparticles encapsulating functional m1Ψ-modified PLPP3 mRNA (0.8 mg/kg). Two weeks post-ligation, carotid arteries and plasma samples were collected. Cross-sections of the ligated LCA were analyzed for lesion size using Oil Red O staining, with plaque areas quantified using ImageJ software. Plasma total cholesterol levels were measured to evaluate systemic lipid profiles.

#### Therapeutic effectiveness of PLPP3 mRNA-encapsulated VCAM1-targeting PROGRAMMED NP in promoting regression of advanced atherosclerotic plaques in mice

A mouse model simulating lesion regression under aggressive cholesterol-lowering therapy, previously described (*5*), was adapted in this study. Briefly, adult male C57BL/6 mice were injected with recombinant AAV serotype-9 expressing mutant PCSK9 under the hepatic control region-apolipoprotein enhancer/alpha1-antitrypsin promoter [AAV9-HCRApoE/hAAT-D377Y-mPCSK9; 1 × 10¹¹ viral genomes (VG), Vector Biolabs] and fed a high-fat diet (TD. 88137, Harlan-Envigo, IN, USA) for 16 weeks. Mice were either sacrificed for baseline analysis (baseline group) or transitioned to a chow diet supplemented with the MTP inhibitor Lomitapide (25 mg/kg chow diet, Research Diet Inc.) for 2 weeks prior to sacrifice (regression group). Regression group mice were stratified into three subgroups and administered three intravenous injections of the following treatments: (1) PBS, (2) VCAM1-targeting PROGRAMMED nanoparticles encapsulating mutant m1Ψ-modified PLPP3 mRNA, or (3) VCAM1-targeting PROGRAMMED nanoparticles encapsulating functional m1Ψ-modified PLPP3 mRNA. The injections (0.8 mg mRNA/kg body weight) were administered via the tail vein on days 1, 6, and 11 following initiation of the chow diet and MTP inhibitor treatment (Fig. 5A). Mice were sacrificed two weeks after starting the chow diet and MTP inhibitor treatment. Blood samples and tissues were collected for analysis. Plasma cholesterol levels were measured to confirm the cholesterol-lowering effect of the chow diet and MTP inhibitor. Hearts were fixed in 4% paraformaldehyde, immersed in 30% sucrose, and embedded in OCT. Serial cross-sections (8 μm thick) of the aortic root were prepared. Oil Red O staining was performed to quantify plaque area. Macrophage accumulation was assessed using CD68 staining (MCA1957, Bio-Rad), while collagen deposition was visualized using Picrosirius red staining. Collagen images were analyzed under a polarizing light microscope, and all data were quantified using ImageJ software.

## Notes

### Competing Interest Statement

The authors have declared no competing interest.

## References

1. M. A. Matthay, R. L. Zemans, G. A. Zimmerman, Y. M. Arabi, J. R. Beitler, A. Mercat, M. Herridge, A. G. Randolph, C. S. Calfee, Acute respiratory distress syndrome. Nat. Rev. Dis. Primers 5, 18 (2019).

2. A. Bonaventura, A. Vecchié, L. Dagna, K. Martinod, D. L. Dixon, B. W. Van Tassell, F. Dentali, F. Montecucco, S. Massberg, M. Levi, Endothelial dysfunction and immunothrombosis as key pathogenic mechanisms in COVID-19. Nat. Rev. Immunol. 21, 319–329 (2021).

3. J. L. Björkegren, A. J. Lusis, Atherosclerosis: recent developments. Cell 185, 1630–1645 (2022).

4. K. Karikó, H. Muramatsu, F. A. Welsh, J. Ludwig, H. Kato, S. Akira, D. Weissman, Incorporation of pseudouridine into mRNA yields superior nonimmunogenic vector with increased translational capacity and biological stability. Mol. Ther. 16, 1833–1840 (2008).

5. D. M. Altmann, R. J. Boyton, COVID-19 vaccination: The road ahead. Science 375, 1127–1132 (2022).

6. O. Soehnlein, P. Libby, Targeting inflammation in atherosclerosis—from experimental insights to the clinic. Nat. Rev. Drug Discov. 20, 589–610 (2021).

7. N. R. London, W. Zhu, F. A. Bozza, M. C. Smith, D. M. Greif, L. K. Sorensen, L. Chen, Y. Kaminoh, A. C. Chan, S. F. Passi, Targeting Robo4-dependent Slit signaling to survive the cytokine storm in sepsis and influenza. Sci. Transl. Med. 2, 23ra19–23ra19 (2010).

8. J. R. Teijaro, K. B. Walsh, S. Cahalan, D. M. Fremgen, E. Roberts, F. Scott, E. Martinborough, R. Peach, M. B. Oldstone, H. Rosen, Endothelial cells are central orchestrators of cytokine amplification during influenza virus infection. Cell 146, 980–991 (2011).

9. E. A. King, J. W. Davis, J. F. Degner, Are drug targets with genetic support twice as likely to be approved? Revised estimates of the impact of genetic support for drug mechanisms on the probability of drug approval. PLoS Genet. 15, e1008489 (2019).

10. M. R. Nelson, H. Tipney, J. L. Painter, J. Shen, P. Nicoletti, Y. Shen, A. Floratos, P. C. Sham, M. J. Li, J. Wang, The support of human genetic evidence for approved drug indications. Nat. Genet. 47, 856–860 (2015).

11. R.-T. Huang, D. Wu, A. Meliton, M.-J. Oh, M. Krause, J. A. Lloyd, R. Nigdelioglu, R. B. Hamanaka, M. K. Jain, A. Birukova, J. P. Kress, K. G. Birukov, G. M. Mutlu, Y. Fang, Experimental lung injury reduces Krüppel-like factor 2 to increase endothelial permeability via regulation of RAPGEF3–Rac1 signaling. Am. J. Respir. Crit. Care Med. 195, 639–651 (2017).

12. S. Xu, Y. Liu, Y. Ding, S. Luo, X. Zheng, X. Wu, Z. Liu, I. Ilyas, S. Chen, S. Han, The zinc finger transcription factor, KLF2, protects against COVID-19 associated endothelial dysfunction. Signal Transduct. Target Ther. 6, 266 (2021).

13. H. Schunkert, I. R. König, S. Kathiresan, M. P. Reilly, T. L. Assimes, H. Holm, M. Preuss, A. F. Stewart, M. Barbalic, C. Gieger, Large-scale association analysis identifies 13 new susceptibility loci for coronary artery disease. Nat. Genet. 43, 333–338 (2011).

14. C. D. Consortium, P. Deloukas, S. Kanoni, C. Willenborg, M. Farrall, T. L. Assimes, J. R. Thompson, E. Ingelsson, D. Saleheen, J. Erdmann, Large-scale association analysis identifies new risk loci for coronary artery disease. Nat. Genet. 45, 25–33 (2013).

15. C. Wu, R.-T. Huang, C.-H. Kuo, S. Kumar, C. W. Kim, Y.-C. Lin, Y.-J. Chen, A. Birukova, K. G. Birukov, N. O. Dulin, M. Civelek, A. J. Lusis, X. Loyer, A. Tedgui, G. Dai, H. Jo, Y. Fang, Mechanosensitive PPAP2B regulates endothelial responses to atherorelevant hemodynamic forces. Circ. Res. 117, 41–53 (2015).

16. M. D. Krause, R.-T. Huang, D. Wu, T.-P. Shentu, D. L. Harrison, M. B. Whalen, L. K. Stolze, A. Di Rienzo, I. P. Moskowitz, M. Civelek, C. E. Romanoski, Y. Fang, Genetic variant at coronary artery disease and ischemic stroke locus 1p32. 2 regulates endothelial responses to hemodynamics. Proc. Natl. Acad. Sci. USA 115, 11349–11358 (2018).

17. G. W. Liu, E. B. Guzman, N. Menon, R. S. Langer, Lipid nanoparticles for nucleic acid delivery to endothelial cells. Pharm. Res. 40, 3–25 (2023).

18. K. J. Kauffman, J. R. Dorkin, J. H. Yang, M. W. Heartlein, F. DeRosa, F. F. Mir, O. S. Fenton, D. G. Anderson, Optimization of lipid nanoparticle formulations for mRNA delivery in vivo with fractional factorial and definitive screening designs. Nano Lett. 15, 7300–7306 (2015).

19. I. Koltover, T. Salditt, J. O. Rädler, C. R. Safinya, An inverted hexagonal phase of cationic liposome-DNA complexes related to DNA release and delivery. Science 281, 78–81 (1998).

20. X. Hou, T. Zaks, R. Langer, Y. Dong, Lipid nanoparticles for mRNA delivery. Nat. Rev. Mater. 6, 1078–1094 (2021).

21. O. F. Khan, E. W. Zaia, H. Yin, R. L. Bogorad, J. M. Pelet, M. J. Webber, I. Zhuang, J. E. Dahlman, R. Langer, D. G. Anderson, Ionizable amphiphilic dendrimer-based nanomaterials with alkyl-chain-substituted amines for tunable siRNA delivery to the liver endothelium in vivo. Angew. Chem. Int. Ed. 53, 14397–14401 (2014).

22. O. F. Khan, E. W. Zaia, S. Jhunjhunwala, W. Xue, W. Cai, D. S. Yun, C. M. Barnes, J. E. Dahlman, Y. Dong, J. M. Pelet, Dendrimer-inspired nanomaterials for the in vivo delivery of siRNA to lung vasculature. Nano Lett. 15, 3008–3016 (2015).

23. Y.-X. Lin, Y. Wang, J. Ding, A. Jiang, J. Wang, M. Yu, S. Blake, S. Liu, C. J. Bieberich, O. C. Farokhzad, Reactivation of the tumor suppressor PTEN by mRNA nanoparticles enhances antitumor immunity in preclinical models. Sci. Transl. Med. 13, eaba9772 (2021).

24. G. Blume, G. Cevc, Liposomes for the sustained drug release in vivo. Biochim. Biophys. Acta Biomembr. 1029, 91–97 (1990).

25. T. Allen, C. Hansen, F. Martin, C. Redemann, A. Yau-Young, Liposomes containing synthetic lipid derivatives of poly (ethylene glycol) show prolonged circulation half-lives in vivo. Biochim. Biophys. Acta Biomembr. 1066, 29–36 (1991).

26. T. M. Allen, P. R. Cullis, Drug delivery systems: entering the mainstream. Science 303, 1818–1822 (2004).

27. J. S. Suk, Q. Xu, N. Kim, J. Hanes, L. M. Ensign, PEGylation as a strategy for improving nanoparticle-based drug and gene delivery. Adv. Drug Del. Rev. 99, 28–51 (2016).

28. Z. Lin, A. Kumar, S. SenBanerjee, K. Staniszewski, K. Parmar, D. E. Vaughan, M. A. Gimbrone Jr, V. Balasubramanian, G. García-Cardeña, M. K. Jain, Kruppel-like factor 2 (KLF2) regulates endothelial thrombotic function. Circ. Res. 96, e48–e57 (2005).

29. K. M. Parmar, H. B. Larman, G. Dai, Y. Zhang, E. T. Wang, S. N. Moorthy, J. R. Kratz, Z. Lin, M. K. Jain, M. A. Gimbrone Jr, Integration of flow-dependent endothelial phenotypes by Kruppel-like factor 2. J. Clin. Invest. 116, 49–58 (2006).

30. K. J. Hassett, K. E. Benenato, E. Jacquinet, A. Lee, A. Woods, O. Yuzhakov, S. Himansu, J. Deterling, B. M. Geilich, T. Ketova, Optimization of lipid nanoparticles for intramuscular administration of mRNA vaccines. Mol. Ther. Nucleic Acids 15, 1–11 (2019).

31. Y. Nakashima, E. W. Raines, A. S. Plump, J. L. Breslow, R. Ross, Upregulation of VCAM-1 and ICAM-1 at atherosclerosis-prone sites on the endothelium in the ApoE-deficient mouse. Arterioscler. Thromb. Vasc. Biol. 18, 842–851 (1998).

32. M. I. Cybulsky, K. Iiyama, H. Li, S. Zhu, M. Chen, M. Iiyama, V. Davis, J.-C. Gutierrez-Ramos, P. W. Connelly, D. S. Milstone, A major role for VCAM-1, but not ICAM-1, in early atherosclerosis. J. Clin. Invest. 107, 1255–1262 (2001).

33. Z. Zhou, C.-F. Yeh, M. Mellas, M.-J. Oh, J. Zhu, J. Li, R.-T. Huang, D. L. Harrison, T.-P. Shentu, D. Wu, M. Lueckheide, L. Carver, E. J. Chung, L. Leon, K.-C. Yang, M. V. Tirrell, Y. Fang, Targeted polyelectrolyte complex micelles treat vascular complications in vivo. Proc. Natl. Acad. Sci. USA 118, e2114842118 (2021).

34. M. Nahrendorf, F. A. Jaffer, K. A. Kelly, D. E. Sosnovik, E. Aikawa, P. Libby, R. Weissleder, Noninvasive vascular cell adhesion molecule-1 imaging identifies inflammatory activation of cells in atherosclerosis. Circulation 114, 1504–1511 (2006).

35. B. R. Anderson, H. Muramatsu, S. R. Nallagatla, P. C. Bevilacqua, L. H. Sansing, D. Weissman, K. Karikó, Incorporation of pseudouridine into mRNA enhances translation by diminishing PKR activation. Nucleic Acids Res. 38, 5884–5892 (2010).

36. D. Wu, T.-H. Lee, R.-T. Huang, R. D. Guzy, N. Schoettler, A. Adegunsoye, J. Mueller, A. Husain, A. I. Sperling, G. M. Mutlu, Y. Fang, SARS-CoV-2 Infection is associated with reduced Krüppel-like factor 2 in human lung autopsy. Am. J. Respir. Cell Mol. Biol. 65, 222–226 (2021).

37. G. Matute-Bello, G. Downey, B. B. Moore, S. D. Groshong, M. A. Matthay, A. S. Slutsky, W. M. Kuebler, An official American Thoracic Society workshop report: features and measurements of experimental acute lung injury in animals. Am. J. Respir. Cell Mol. Biol. 44, 725–738 (2011).

38. L. Guo, Y.-C. Wang, J.-J. Mei, R.-T. Ning, J.-J. Wang, J.-Q. Li, X. Wang, H.-W. Zheng, H.-T. Fan, L.-D. Liu, Pulmonary immune cells and inflammatory cytokine dysregulation are associated with mortality of IL-1R1-/-mice infected with influenza virus (H1N1). Zool. Res. 38, 146 (2017).

39. L. P. Dawson, M. Lum, N. Nerleker, S. J. Nicholls, J. Layland, Coronary atherosclerotic plaque regression: JACC state-of-the-art review. J. Am. Coll. Cardiol. 79, 66–82 (2022).

40. A. Sarraju, S. E. Nissen, Atherosclerotic plaque stabilization and regression: a review of clinical evidence. Nat. Rev. Cardiol., 1–11 (2024).

41. S. E. Nissen, S. J. Nicholls, I. Sipahi, P. Libby, J. S. Raichlen, C. M. Ballantyne, J. Davignon, R. Erbel, J. C. Fruchart, J.-C. Tardif, Effect of very high-intensity statin therapy on regression of coronary atherosclerosis: the ASTEROID trial. JAMA 295, 1556–1565 (2006).

42. B. Hewing, S. Parathath, C. K. Mai, M. I. Fiel, L. Guo, E. A. Fisher, Rapid regression of atherosclerosis with MTP inhibitor treatment. Atherosclerosis 227, 125–129 (2013).

43. M. Peled, H. Nishi, A. Weinstock, T. J. Barrett, F. Zhou, A. Quezada, E. A. Fisher, A wild-type mouse-based model for the regression of inflammation in atherosclerosis. PLoS One 12, e0173975 (2017).

44. S. S. Martin, A. W. Aday, Z. I. Almarzooq, C. A. Anderson, P. Arora, C. L. Avery, C. M. Baker-Smith, B. Barone Gibbs, A. Z. Beaton, A. K. Boehme, 2024 heart disease and stroke statistics: a report of US and global data from the American Heart Association. Circulation 149, e347–e913 (2024).

45. K. Koelle, M. A. Martin, R. Antia, B. Lopman, N. E. Dean, The changing epidemiology of SARS-CoV-2. Science 375, 1116–1121 (2022).

46. D. A. Berlin, R. M. Gulick, F. J. Martinez, Severe covid-19. N. Engl. J. Med. 383, 2451–2460 (2020).

47. P. Libby, The changing landscape of atherosclerosis. Nature 592, 524–533 (2021).

48. M. J. Henley, A. N. Koehler, Advances in targeting ‘undruggable’transcription factors with small molecules. Nat. Rev. Drug Discov. 20, 669–688 (2021).

49. B. A. Smith, C. R. Bertozzi, The clinical impact of glycobiology: targeting selectins, Siglecs and mammalian glycans. Nat. Rev. Drug Discov. 20, 217–243 (2021).

50. A. S. Slutsky, V. M. Ranieri, Ventilator-induced lung injury. N. Engl. J. Med. 369, 2126–2136 (2013).

51. G. B. Mills, W. H. Moolenaar, The emerging role of lysophosphatidic acid in cancer. Nat. Rev. Cancer 3, 582–591 (2003).

52. T. Hla, M.-J. Lee, N. Ancellin, J. H. Paik, M. J. Kluk, Lysophospholipids--receptor revelations. Science 294, 1875–1878 (2001).

53. P. M. Ridker, B. M. Everett, T. Thuren, J. G. MacFadyen, W. H. Chang, C. Ballantyne, F. Fonseca, J. Nicolau, W. Koenig, S. D. Anker, Antiinflammatory therapy with canakinumab for atherosclerotic disease. N. Engl. J. Med. 377, 1119–1131 (2017).

54. J. C. Cohen, H. H. Hobbs, Simple genetics for a complex disease. Science 340, 689–690 (2013).

55. R. S. Rosenson, R. A. Hegele, S. Fazio, C. P. Cannon, The evolving future of PCSK9 inhibitors. J. Am. Coll. Cardiol. 72, 314–329 (2018).

56. I. Hekselman, E. Yeger-Lotem, Mechanisms of tissue and cell-type specificity in heritable traits and diseases. Nat. Rev. Genet. 21, 137–150 (2020).

57. K. A. Jagadeesh, K. K. Dey, D. T. Montoro, R. Mohan, S. Gazal, J. M. Engreitz, R. J. Xavier, A. L. Price, A. Regev, Identifying disease-critical cell types and cellular processes by integrating single-cell RNA-sequencing and human genetics. Nat. Genet. 54, 1479–1492 (2022).

58. G. R. Schnitzler, H. Kang, S. Fang, R. S. Angom, V. S. Lee-Kim, X. R. Ma, R. Zhou, T. Zeng, K. Guo, M. S. Taylor, Convergence of coronary artery disease genes onto endothelial cell programs. Nature 626, 799–807 (2024).

## References for the Supplementary Materials and Methods

1. G. Matute-Bello, G. Downey, B. B. Moore, S. D. Groshong, M. A. Matthay, A. S. Slutsky, W. M. Kuebler, An official American Thoracic Society workshop report: features and measurements of experimental acute lung injury in animals. Am. J. Respir. Cell Mol. Biol. 44, 725–738 (2011).

2. L. Guo, Y.-C. Wang, J.-J. Mei, R.-T. Ning, J.-J. Wang, J.-Q. Li, X. Wang, H.-W. Zheng, H.-T. Fan, L.-D. Liu, Pulmonary immune cells and inflammatory cytokine dysregulation are associated with mortality of IL-1R1-/-mice infected with influenza virus (H1N1). Zool. Res. 38, 146 (2017).

3. D. Nam, C.-W. Ni, A. Rezvan, J. Suo, K. Budzyn, A. Llanos, D. Harrison, D. Giddens, H. Jo, Partial carotid ligation is a model of acutely induced disturbed flow, leading to rapid endothelial dysfunction and atherosclerosis. Am. J. Physiol. Heart Circ. Physiol. 297, 1535–1543 (2009).

4. C. Wu, R.-T. Huang, C.-H. Kuo, S. Kumar, C. W. Kim, Y.-C. Lin, Y.-J. Chen, A. Birukova, K. G. Birukov, N. O. Dulin, M. Civelek, A. J. Lusis, X. Loyer, A. Tedgui, G. Dai, H. Jo, Y. Fang, Mechanosensitive PPAP2B regulates endothelial responses to atherorelevant hemodynamic forces. Circ. Res. 117, e41–e53 (2015).

5. M. Peled, H. Nishi, A. Weinstock, T. J. Barrett, F. Zhou, A. Quezada, E. A. Fisher, A wild-type mouse-based model for the regression of inflammation in atherosclerosis. PLoS One 12, e0173975 (2017).

